# A hypoxia-sensitive medullary nucleus modulates cerebral blood flow via a disynaptic pathway to cortex

**DOI:** 10.64898/2026.02.20.707130

**Authors:** Karishma Chhabria, Jacob Duckworth, Pantong Yao, Arash Fassihi-Zakeri, David Kleinfeld

## Abstract

The RVLM (rostral ventral lateral medulla) region of the brainstem is implicated as a controller of both systemic and cerebral blood flow (CBF). Past studies leave open the question of how the RVLM stabilizes CBF in awake animals. Here, we focus on CBF regulation by the adenergic subpopulation of RVLM neurons (RVLM_Dβh_). Hypoxic challenge increases the variability of the activity of RVLM_Dβh_ neurons along with increased CBF. Experimental photoactivation of RVLM_Dβh_ neurons leads to rapid vasodilation of pial arterioles across the cortical mantle and increased CBF, which is only then followed by an increase in cortical activity. No significant changes in systemic physiology are observed. Virus tracing establishes disynaptic pathways from the RVLM to neocortex with predominant relays involving the lateral hypothalamus and the zona incerta subthalamic nuclei. Chemogenetic inhibition of those nuclei led to a 70 % reduction in the ability of photoactivated RVLM_Dβh_ neurons to induce cortical vasodilation and increase CBF. In toto, these findings reveal a major RVLM subcortical pathway to drive transcortical increases in CBF and represent an adaptive mechanism to inform the cerebral vasculature about environmental shifts in pO_2_.

## INTRODUCTION

Cerebral blood flow (CBF) is a tightly regulated physiological process, ensuring that the brain’s energetic demands are met across changing environments. Small disruptions in perfusion or an inadequate response to changes in environmental oxygen (pO_2_) can have catastrophic physiological and behavioral outcomes. Multiple mechanisms act in concert to regulate CBF across spatial and temporal scales [1-3]. Slowly activating mechanisms of CBF control include those mediated by modulatory transmitters [4], notably, prostaglandins [5] and adenosine [6], and by pericyte activity [7]. In contrast, fast mechanisms involve propagation of potassium action potentials from microvessels to upstream arterioles via gap junctions between endothelial cells and between endothelial and smooth muscle cells [8]. Fast mechanisms may also involve specific types of cortical neurons [9-12] and direct synaptic contacts [13]. Beyond these local control mechanisms, neuromodulators released from subcortical nuclei have been shown to alter cerebrovascular tone [14-16]. However, it remains unknown how the brain dynamically adjusts perfusion in response to physiological challenges such as changes in the ambient gas composition that accompany activities in awake animals, such as diving, high-altitude exposure, or intense physical exertion, to ensure functioning of central circuits. Understanding this coupling between cerebrovascular control and physiological context is essential to reveal how the brain maintains homeostasis under fluctuating systemic demands.

A bridge between changes in systemic physiology and brain blood flow involves the rostral ventrolateral medulla (RVLM), a part of baroreflex circuitry [17, 18] that helps maintains peripheral vascular resistance and hence blood pressure during systemic challenges [19, 20]. In contrast, much less is known about the central adjustments in CBF required to maintain systemic homeostasis. Pioneering functional studies by Reis and Golanov [21] demonstrated that electrical stimulation of the RVLM can modulate CBF in both intact and in spinalized anesthetized rats. Yet anesthesia markedly affects the RVLM’s influence on the peripheral vasculature [22]. Given that neurons in the RVLM form ascending intracerebral projections, along with projections to the spinal intermediolateral column [23], it is imperative to establish the RVLM’s role in the control of CBF in awake animals.

The heterogeneity of cell types that comprise the RVLM include adrenergic [24], glutamatergic [25, 26], GABAergic [27], and glycinergic [28] expressing neurons. This adds complexity to the interpretation of experiments. An important advance to unravel RVLM function is highlighted by the work of Card and colleagues [29] with their demonstration that the adrenergic neuron subpopulation can be selectively identified from their expression of dopamine beta hydroxylase (Dβh) in transgenic mice. Furthermore, these adrenergic neurons, denoted by RVLM_Dβh_ neurons, make up a predominant portion of the RVLM [30].

Here we ask: (*i*) In awake mice, what is the nature of RVLM_Dβh_ activation in response to hypoxia? We are motivated in part by a lack of studies on the neuronal response to hypoxia other than by barosensitive bulbospinal neurons, as found in vitro using whole-cell patch clamp recordings [31]. (*ii*) Can selective activation of the RVLM_Dβh_ neurons modulate cortical vascular networks and thus CBF? Global, electrical activation of the RVLM, which likely activates all neuronal subtypes, will robustly modulate CBF [32]. However, the specific contribution of RVLM_Dβh_ neuronal activity to CBF is unknown. Even in the more heavily-studied case of systemic vascular regulation, the contribution of RVLM_Dβh_ neurons is controversial [22], with evidence suggesting that non-adrenergic bulbospinal neuronal populations strongly contribute to the control of peripheral vascular tone. (*iii*) What is the ascending circuitry that underlies the influence of RVLM_Dβh_ neurons on CBF modulation? Pioneering studies used electrical measurements and excitotoxicity blocks to infer relays from the RVLM to cortex, and identified a putative trisynaptic pathway [21]. Here, we use the state-of-the-art reverse engineering tools of viral labeling, optogenetic activation, and chemogenic transient blocks of activity in awake animals [33] to guide our search for potential direct connections as well as disynaptic pathways from RVLM_Dβh_ neurons to cortical somata.

## RESULTS

### Hypoxia increases variability in the activity of RVLM_Dβh_ neurons

We first assessed the potential correlation between activity of RVLM_Dβh_ neurons and changes in CBF, as measured by laser Doppler flowmetry (LDF), under low ambient oxygen. The RVLM region of Dβh-Cre mice were injected with AAV-Flex-GCaMP8s (**Figure 1A**) and implanted with an optic fiber opposing the surface of the RVLM. These mice were placed in a custom flow chamber that enabled us to modulate the flow of oxygen (**Figure 1B,C**), We tested if the intracellular calcium concentration in RVLM_Dβh_ neurons, denoted [Ca^2+^], correlates with the changes in CBF. As a control for neural consequences of hypoxia, we observed pupil constriction (**Figure 1D**), consistent with a decrease in arousal [34]. We further observed that hypoxia leads to an expected increase in the CBF (**Figure 1E**). The new result is an increase in the variability of [Ca^2+^] in RVLM_Dβh_ neurons, albeit not the mean, in response to hypoxia (**Figure 1F**).

**Figure 1.**
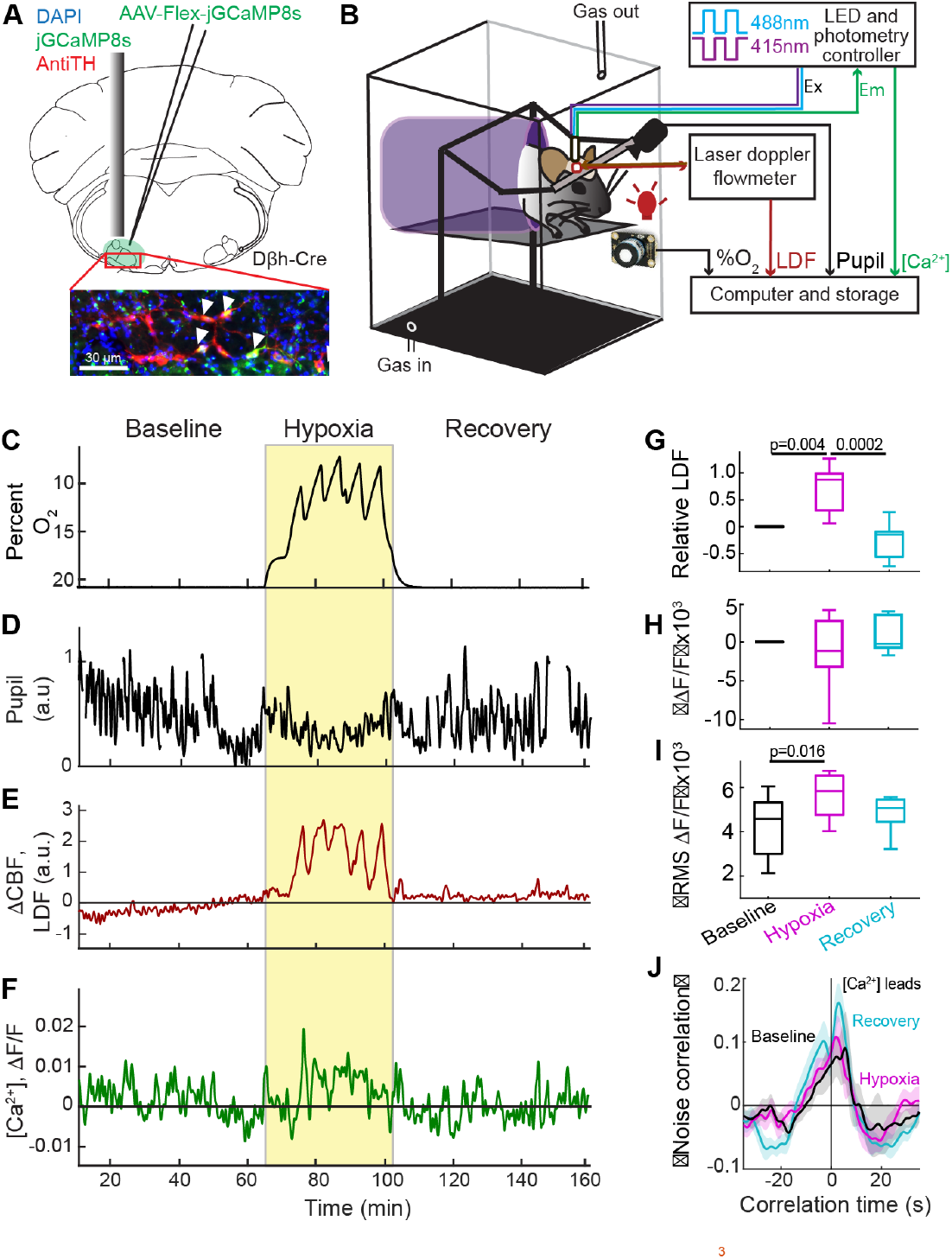
Hypoxia responses of RVLM_Dβh_ neurons and CBF in awake head fixed micex. (**A**) Injection cartoon showing injection site of AAV9-CAG-Flex-jGCaMP8s-WPRE-SV40, fiber implantation and cells labelled at injection site. Scale bar: 30 μm. (**B**) Schematic showing the hypoxia chamber set up with the hammock, gas inlets/outlets, oxygen sensor, infrared LED, and camera for pupillometry. Fiber photometry signals and cortical CBF are measured from an awake, head-fixed mice. (**C**) Oxygen (O_2_) sensor reading throughout one hypoxia trial showing percent change in O_2_. Gas flow was from 0 to 160 minutes. (**D**) Pupil diameter recorded throughout the hypoxia trial showing decrease in arousal during hypoxia. (**E**) Smoothed (zero phase Savitzky-Golay filter) LDF signal recorded simultaneously from the parietal cortex in awake mouse showing increase during hypoxia. (**F**) Smoothed (zero phase Savitzky-Golay filter) time series of ΔF/F calculated from calcium [Ca^2+^] signal recorded from RVLM_Dβh_ neurons for the hypoxia trial. The mean LDF in the different regions are: 0 (baseline), 0.78 (hypoxia) and 0.25 (recovery). The mean and root mean square (RMS) in different regions is: 0 and 0.004 (baseline); -0.001 and 0.006 (hypoxia); 0.001 and 0.005 (recovery). (**G**) Relative LDF calculated as change from baseline for 7 mice showing hypoxia significantly increases CBF compared to baseline (p = 0.004, paired t-test) and recovery (p = 0.002). (**H**) Mean ΔF/F, across 7 mice showing no significant change during hypoxia compared to baseline (p = 0.53) and recovery (p = 0.25). (**I**) RMS of ΔF/F across 7 mice showing a significant increase during hypoxia compared to baseline (p = 0.016, Wilcoxon test) however not significant when compared to recovery (p = 0.46). (**J**) Mean correlation of the [Ca^2+^] from RVLM_Dβh_ neurons with the CBF signals showing [Ca^2+^] leads CBF by 2.94 s during baseline, 1.94 s and 5.46 s during hypoxia and recovery (7 mice).

As a summary across animals (7 mice), the CBF is significantly increased compared to baseline and recovery (p *=* 0.001, repeated measures ANOVA) (**Figure 1G**). The mean of the calcium signal in RVLM_Dβh_ neurons in unchanged (**Figure 1H**). Yet the variance of the calcium signal in RVLM_Dβh_ neurons, characterized by the change in the root mean square (RMS) [Ca^2+^] in RVLM_Dβh_ neurons during hypoxia significantly increased relative to baseline (p *=* 0.019, Wilcoxon’s test) (**Figure 1I**). It is important to recall that driving an increase in downstream smooth muscle activity, like the case of neural signaling, can involve changes in variability as well as changes in mean activity [35]. Lastly, we calculated the noise correlation between the CBF and the change in calcium signal in RVLM_Dβh_ neurons. This addresses the timing of potential control of CBF by the RVLM. The calcium signal leads the LDF signal before, during, and after hypoxia, implying that the timing of RVLM_Dβh_ signaling is maintained during hypoxia while hypoxic increases in RVLM_Dβh_ variability affects the amplitude (**Figure 1J**).

### Activation of RVLM_Dβh_ neurons dilates cortical arterioles

To go beyond the correlation between the activity of RVLM_Dβh_ neurons and CBF (**Figure 1**), we tested whether RVLM_Dβh_ neurons can exert a causal influence on CBF. To this end, we infected Dβh-Cre mice with an AAV virus that was constructed to express channelrhodopsin (ChR) in RVLM_Dβh_ neurons (**Figure 2A**). We used these mice to check if optogenetic stimulation of the RVLM_Dβh_ neurons led to an increase in the diameter of cortical arterioles. These measurements were performed by imaging individual pial arterioles in parietal cortex with adaptive optics / two-photon microscopy [36] concurrent with photostimulation of RVLM_Dβh_ neurons through an implanted fiber (**Figure 2B,C**). Noting that previous in vitro studies showed that RVLM neurons fire over a broad range of frequencies [37, 38], we tested the response to a single 5 s pulse with a modulated intensity, i.e., 10, 20, and 50 Hz, versus continuous intensity for *in vivo* optogenetic stimulation (**Figure 2B,C**; **Extended Figure 2C**,**D**). We selected continuous illumination as most effective single pulse (**Figure 2C**).

**Figure 2.**
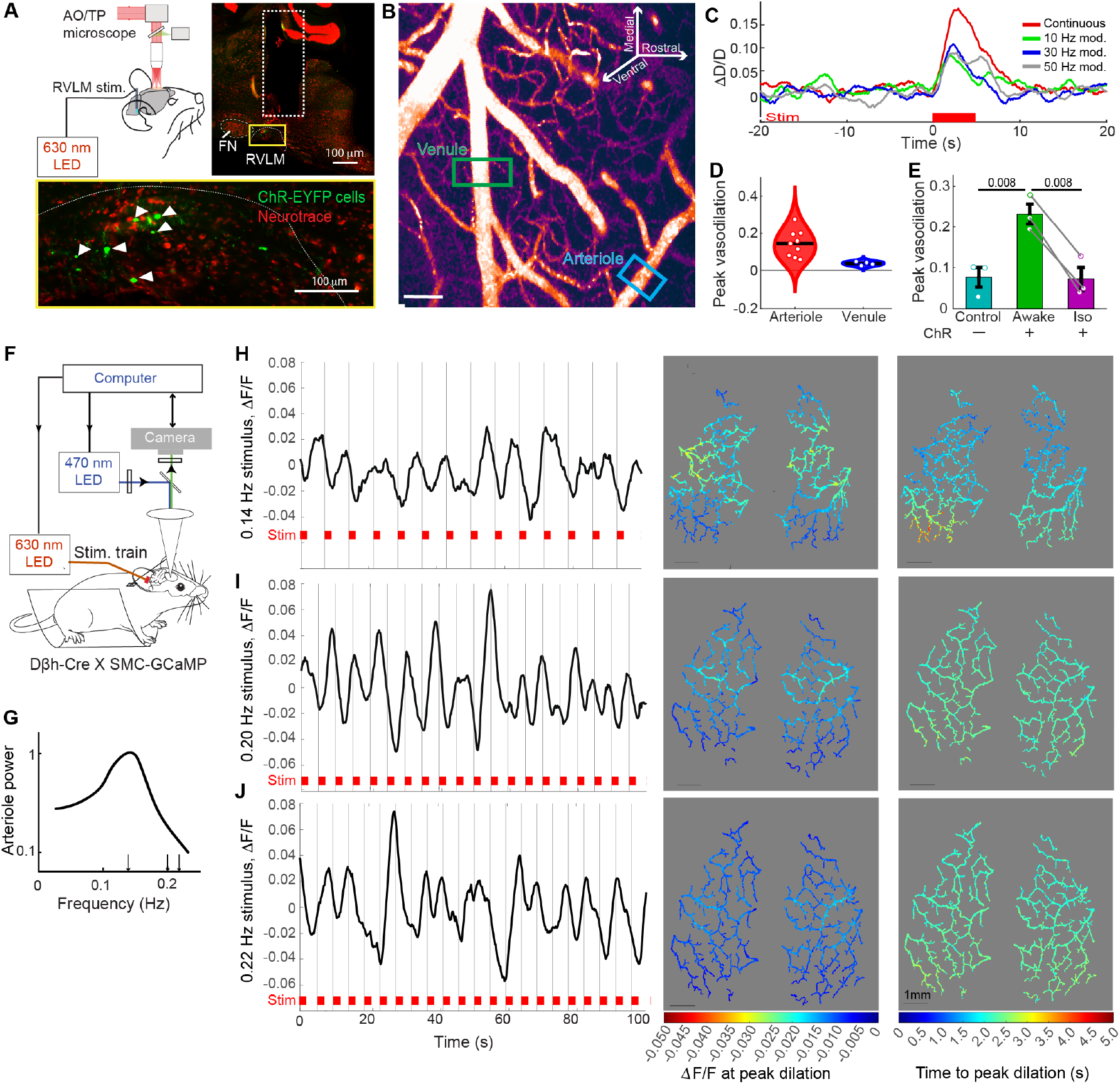
Activation of RVLM_Dβh_ neurons increases CBF by dilating cortical arterioles. (**A**) Schematic showing location of injection of AAVDJ-EF1α-DIO-hChR2(H134R)-eYFP-WPRE-pA, optic fiber location for optogenetic stimulation of RVLM_Dβh_ neurons and adaptive optics / two-photon microscope. (**B**) Image of brain surface with AO/2P microscopy. Scale bar: 50 μm. The boxes indicate vessels under observation in response to 5 s stimulation. (**C**) Single trial showing vasodilation expressed as change in diameter (ΔD/D) with respect to the baseline in response to 5 s continuous pulse, 10, 30, or 50 Hz stimulation. (**D**) Peak vasodilation (peak of ΔD/D) of pial arterioles and venules in response to a 5 s stimulation showing fractional increase of 0.15 in arterioles (9 arteries across 4 mice) and 0.04 in venules (4 veins in 4 mice). (**E**) Peak vasodilation of pial arterioles showing peak vasodilation of 0.15 (3 arterioles in 3 mice) in awake ChR expressing mice. This response was significant (p = 0.008, unpaired t-test) when compared vasodilation of 0.08 in mice expressing EYFP only (3 mice). One percent (v/v) Isoflurane significantly reduced the vasodilation to 0.07 in mice expressing ChR (p = 0.008, paired t-test, 3 mice). (**F**) Schematic of widefield recording of cortical smooth muscle-cell calcium [Ca^2+^]_i_ with simultaneous stimulation of RVLM_Dβh_ neurons expressing AAV8-EF1α-DIO-ChRmine-eYFP-WPRE using an LED placed in the auditory canal in Dβh-Cre X Tg(Acta2-GCaMP8.1/mVermilion)B34-4Mik/J. (**G**) Typical pial arteriole spectral activity. Note the sharp drop in response for frequencies above 0.2 Hz. (**H-J**) Smooth muscle [Ca^2+^]_i_, expressed as ΔF/F and averaged over the whole cortex for 2 s-wide pulses of 50 Hz optogenetic stimulation, presented at 0.14, 0.20, and 0.22 Hz. Corresponding spatial maps showing peak hyperpolarization responses (peak -ΔF/F) and time-to-peak vasodilation(s) in different vessel segments. Scale bar: 1 mm.

The magnitude of arteriole dilation following excitation to RVLM_Dβh_ neurons was variable but consistently higher in arterioles than venules (9 arterioles, 4 venules across 4 mice) (**Figure 2D, Extended Figure 2A**,**B**). The arteriole dilation is negligible in control mice and was reduced under isoflurane anesthesia (3 mice, p = 0.035; paired t-test) (**Figure 2E**). These data support a causal role for RVLM in the modulation of CBF at the level of single pial arterioles in cortex.

Our next measurements concerned the steady-state spatial extent of RVLM control over vasodilation of cortical arterioles. We performed widefield imaging of pial arterioles in Dβh-Cre mice that were crossed to expressed GCaMP8.1 in vascular smooth-muscle cells [39]. Further, the RVLM in these mice were injected with AAV containing a red-shifted opsin, ChRmine [40], to enable optogenetic stimulation of RVLM_Dβh_ neurons (**Figure 2F**). In these experiments, the arteriole [Ca^2+^]_i_ signal is a surrogate for arteriole diameter; decreases in [Ca^2+^]_i_ correspond to dilation, and vice versa [39]. The periodic optogenetic stimulation of RVLM_Dβh_ neurons used three stimulus frequencies for a periodic train of pulses (**Figure 2G)**. One train was close to the vasomotor peak, i.e., 0.14 Hz (**Figure 2H**), and the other trains at higher frequencies, i.e., 0.20 and 0.22 Hz (**Figure 2I,J**), where a reduced response is expected based on the vasodynamics of pial arterioles [39].

We observed that the spatiotemporal decrease in smooth-muscle [Ca^2+^] emerged most strongly and most rapidly, i.e., within ∼ 2 s of the stimulus onset, for stimulation close to the vasomotor peak (**Figure 2H**). The rostral to caudal spatial gradient across parietal cortex is like that observed in the resting state [41]. All told, these findings demonstrate that activation of RVLM_Dβh_ neurons increases CBF via dilation of arterioles across the cortical mantle.

### Activation of RVLM_Dβh_ neurons increases CBF prior to heightened cortical neuronal activity

Absent stimulation of RVLM, both ongoing and stimulus induced neuronal activity precedes the changes in CBF by ∼ 2 s, measuring from the onset of cortical neural activity until the peak arteriole dilation [42, 43]. To check if this sequence of activation holds for modulation of CBF by activation of the RVLM, we again made use of Dβh-Cre mice that expressed ChR in RVLM_Dβh_ neurons (**Figure 2A**) and concurrently measured both the LDF, to quantify CBF, and the electrocorticogram (ECoG), to measure regional neuronal activity, in response to optogenetic stimulation of RVLM_Dβh_ neurons (**Figure 3A**).

**Figure 3.**
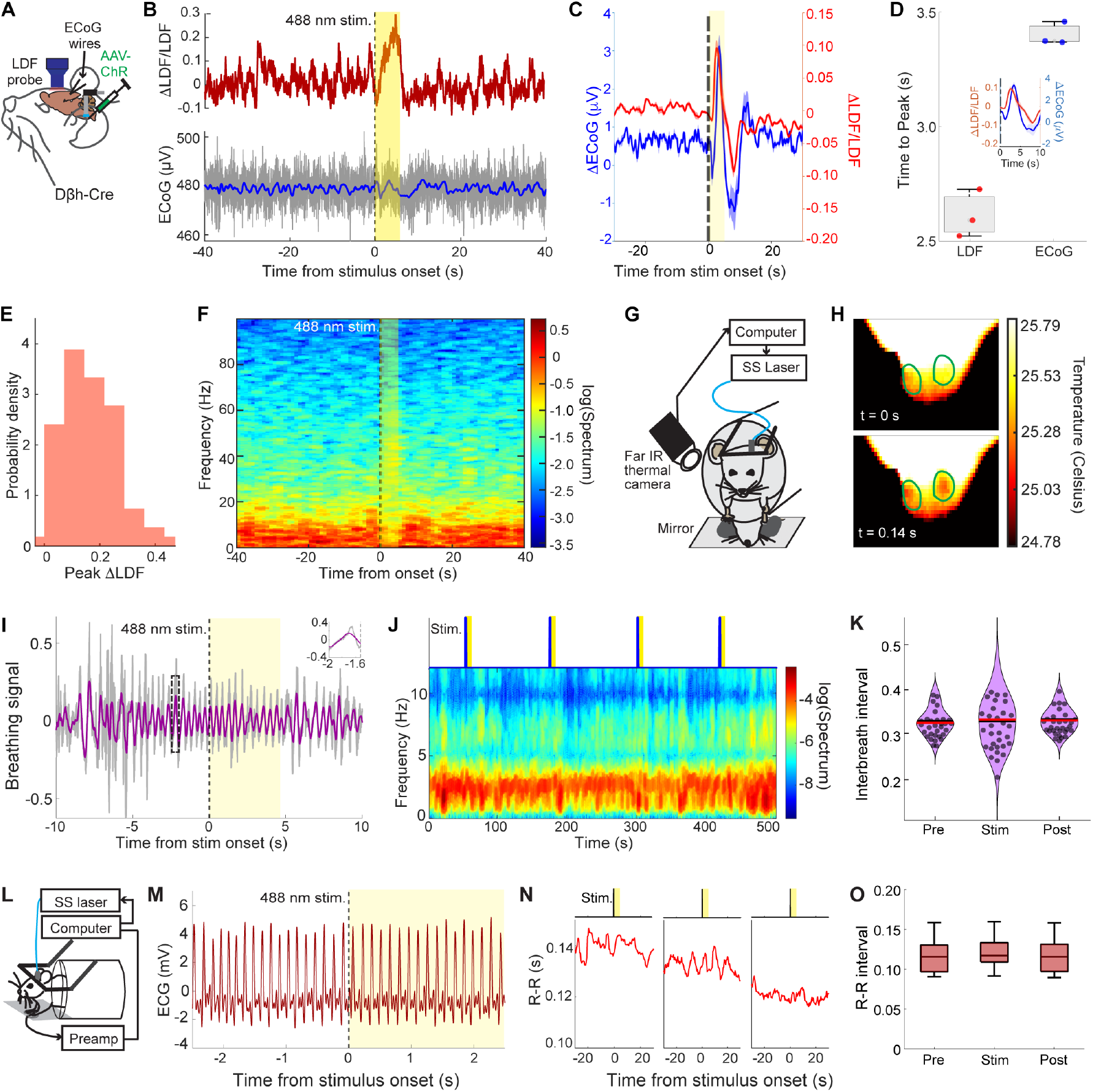
Activation of RVLM_Dβh_ neurons increases CBF with without significantly affecting heart rate and respiration. (**A**) Schematic showing injection site of AAVDJ-EF1α-DIO-hChR2(H134R)-eYFP-WPRE-pA, optic fiber location, electrocorticogram (ECoG) wire placement, and LDF probe placement over thinned parietal cortical window. (**B**) Single trial recorded from an awake mouse showing normalized LDF signal and corresponding ECoG recording from the parietal cortex with 5 s of optogenetic activation of RVLM_Dβh_ neurons (shaded area). The filtered ECoG is shown in blue overlapping the raw signal in gray. (**C**) Mean ECoG and LDF signals from awake mice (3 mice and 75 trials) showing increase in both signals (peak ECoG = 0.35 μV, peak ΔLDF/LDF = 0.09) with 5 s of activation of RVLM_Dβh_ neurons. (**D**) Mean time to peak comparison showing LDF peak (2.68 s from stimulus onset) precedes ECoG (3.45 s from stimulus onset) across 3 mice and 75 trials. Inset shows the time series of mean ECoG and LDF for 10 s form stimulus onset highlighting the difference in time to peak. (**E**) Probability density distribution of peak LDF responses (7 mice and 75 trials) showing 17 % (0.17 ± 0.09) increase in LDF from baseline with stimulation of RVLM_Dβh_ neurons. (**F**) Spectrogram calculated from the ECoG signal recorded in panel B showing negligible changes in the frequency bands with stimulation of RVLM_Dβh_ neurons. Mean spectral power in gamma band (30 to 60 Hz) during prestimulus, stimulus, and post stimulus periods are 0.011, 0.013, and 0.011, respectively. (**G**) Schematic showing breathing recording set up using far infrared (FLIR) thermal camera in awake mouse. (**H**) Images that show changes in the nostril temperature across one inhalation. (**I**) Example trace showing raw (gray) and filtered (purple) breathing signal for one experiment with stimulation of RVLM_Dβh_ neurons. (**J**) Spectrogram of breathing signal illustrating the breathing frequency band (2 to 4 Hz) across multiple RVLM_Dβh_ stimulation pulses showing consistent power in the breathing frequency band. (**K**) Mean interbreath intervals (IBI) calculated from recorded nostril temperature during prestimulus (Pre), stimulus (Stim), and post stimulus (Post) epochs were 0.327, 0.327 and 0.331 s across mice (3 mice, 50 trials, p = 0.84, repeated measures ANOVA) averaged across mice. (**L**) Schematic showing electrocardiogram (ECG) recording, acquisition, and optogenetic stimulation of RVLM_Dβh_ neurons via optical fiber in awake mouse. (**M**) Typical recording showing filtered ECG signal before and during 2.5 s of stimulation. (**N**) R-R intervals calculated over windows of 10 s from recorded ECG across three experiments showing negligible effect of stimulation of RVLM_Dβh_ neurons. (**O**) Mean R-R intervals averaged across Pre, Stim and Post durations were 0.125, 0.128 and 0.129 s, respectively (32 trials across 3 mice; p = 0.70, *repeated measures ANOVA*).

We observed a robust increase in the amplitudes of the CBF and the ECoG in awake mice concurrent with stimulation of RVLM_Dβh_ neurons (**Figure 3B**). The averaged CBF and ECoG traces both exhibited a biphasic response, with an initial rise followed by a transient suppression and return to baseline (**Figure 3B,C**). The change in CBF peaks ∼ 2.5 s after stimulation and, of note, precedes the ECoG peak by ∼ 0.8 s (75 trials across 3 mice) (**Figure 3D**). Across our cohort, stimulation of RVLM_Dβh_ neurons produced a 17 ± 9 percent increase in CBF relative to baseline (**Figure 3E**). The presence of a change in blood flow prior to a change in the electrical activity of cortex provides evidence that the increase in CBF is driven by a mechanism extrinsic to cortex.

A final observation is that the increase in CBF triggered by activation of RVLM_Dβh_ neurons occurs without an increase in gamma-band (30 to 60 Hz) power in the ECoG (**Figure 3F**) (75 trials across 3 mice; p = 0.25). Such an increase is associated with activation of parvalbumin neurons [12, 44-46] and the variation in gamma-band power underlies ∼ 0.1 Hz neural oscillations in the brain [42, 47]. This observation is further suggestive of cortex-wide excitation by the RVLM via a mechanism that is extrinsic to cortex.

### RVLM_Dβh_ neurons modulate CBF without inducing systemic physiological changes

Whole body changes in physiology could influence blood flow dynamics in the brain, much as the mechanisms of cerebral autoregulation purportedly counteract such influence [48, 49]. We thus performed control experiments to ascertain that the increase in CBF signaled by activity of RVLM_Dβh_ neurons was driven by a brain mechanism, as opposed to a change in systemic physiology. First, we monitored breathing with the use of a far infrared thermal camera aimed at the nostrils of head-fixed mice (**Figure 3G-I**). Spectral analysis of the breathing signal revealed a dominant 2 to 4 Hz rhythm that remained unchanged during stimulation (**Figure 3J**). The analysis of the time series of interbreath intervals (IBI) confirmed a lack of significant differences across our cohort (50 trials across 3 mice; p = 0.84) (**Figure 3K**).

We next assessed cardiac output by recording the electrocardiogram (ECG) across the forepaws during optogenetic stimulation of RVLM_Dβh_ neurons in head fixed mice (**Figure 3L**). Distinct P–Q–R complexes, which correspond to depolarization of the right and left ventricles and contraction of the large ventricular muscles, and T waves, which represents repolarization of the ventricles, were reliably detected (**Figure 3M**). Analysis of R–R intervals revealed no significant change in heart rate between baseline and stimulation across our cohort (32 trials across 3 mice; p = 0.70) (**Figure 3N,O**).

These new data on breathing and cardiac output add to prior measurements that showed that activation of RVLM_Dβh_ neurons did not increase systemic blood pressure [50]. All told, the results from optogenetic activation of RVLM_Dβh_ neurons demonstrate that the RVLM can increases CBF prior to, and thus interpreted as independent of, local neuronal activity in the awake animal (**Figures 2** and **3A-F**). Further, the increase in CBF is independent of systemic physiological parameters in the awake animal (**Figure 3G-O**). These findings differ from previous results obtained using electrical stimulation of the RVLM in anesthetized rodents [51], as well as from hypoxia-induced activation of the RVLM, the latter of which produced apnea and bradycardia [52, 53]. By contrast, our results are consistent with the results from studies with anesthetized and spinalized rats, which show increases in CBF following RVLM stimulation even when systemic sympathetic pathways are severed [54].

### Anterograde projections of RVLM_Dβh_ neurons localize in subcortical brain regions

Having established that activation of RVLM_Dβh_ neurons drives cortical arteriole vasodilation, we sought to map the circuitry that underlies this control. We first examined whether projections from RVLM_Dβh_ neurons directly reach cortical somata to form monosynaptic connections. We expressed a *Cre*-dependent anterograde AAV tracer, tagged with mCherry, in Dβh-Cre mice (**Figure 4A**). Animals were permitted to survive for 4 to 5 weeks, after which they were prepared for histology. Sections were stained with fluorescently labeled antibodies to tyrosine hydroxylase (TH) and a fluorescent counterstain and then scanned (**Figure 4A,B**). Somata that expressed mCherry were observed between 6.5 mm and 6.9 mm from bregma post-mortem and co-labeled by TH immunostaining to confirm their adrenergic identity in the RVLM (**Figure 4A,B**). Fibers labeled with mCherry, including axons and fibers of passage, were distributed extensively from the brainstem to the forebrain, consistent with prior studies in rats [29].

**Figure 4.**
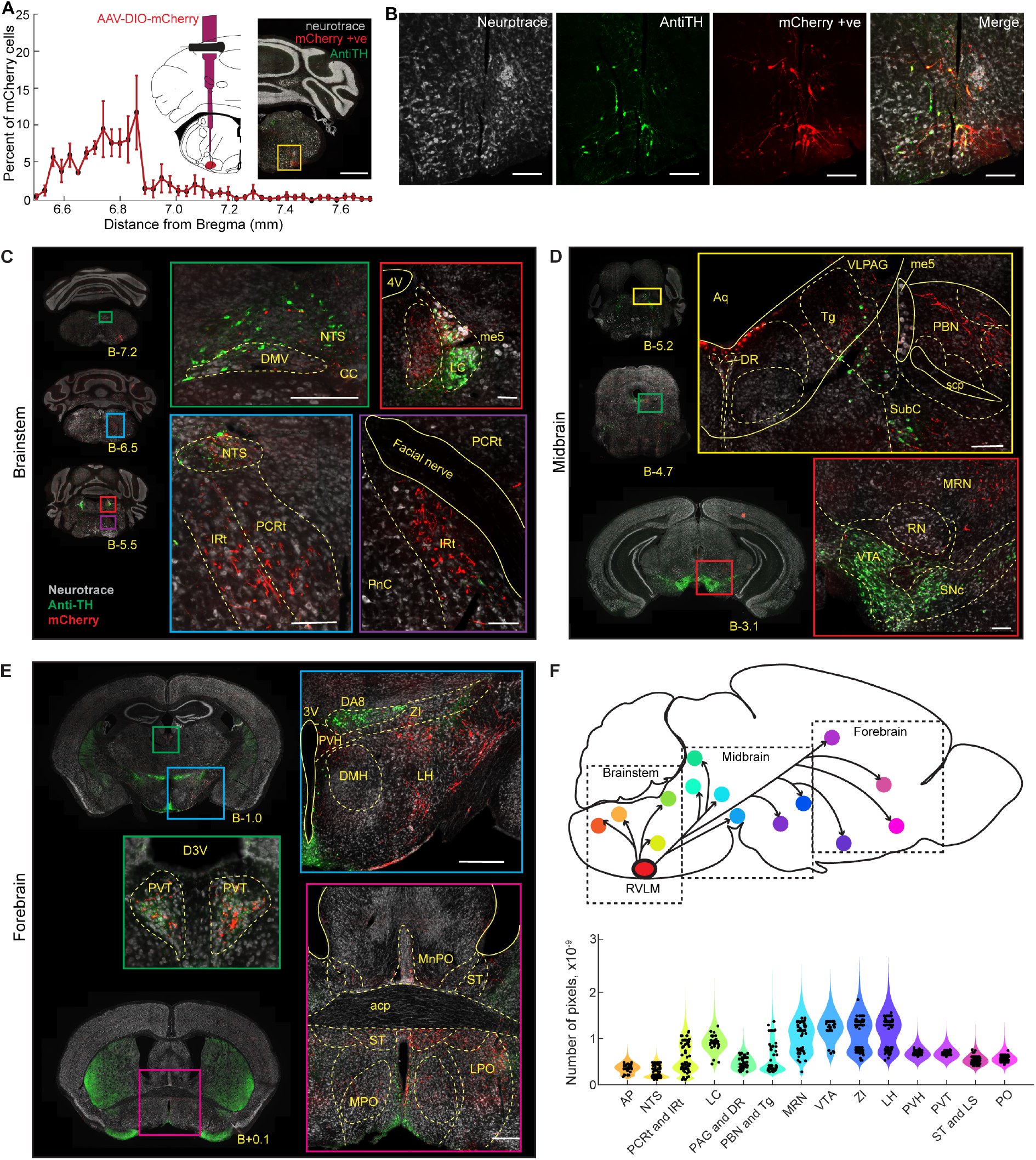
Primary ascending targets of RVLM_Dβh_ neurons across the brain. (**A**) Schematic showing the injection location of AAV8-EF1α-DIO-mCherry and the rostrocaudal extent of labelled cells at the injection site with reference to bregma (B). Scale bar: 100 μm. (**B**) Coronal section showing mCherry positive cells at the injection site, where section was counterstained with Neurotrace blue to localize neuronal somata and Tyrosine hydroxylase (TH) antibody to corroborate adrenergic identity of labelled cells. (**C**) Brainstem labelling of axonal fibers originating from the neuronal somata at the injection site. Fibers were observed in nucleus solitaris (NTS), dorsal motor vagal nucleus (DMV), parvicellular and intermediate parts of reticular nucleus (PCRt and IRt, respectively) and locus coeruleus (LC). (**D**). Midbrain labelling included fibers in venterolateral paraqueductal gray (VLPAG), dorsal raphe (DR), parabrachial nucleus (PBN), tegmentum (Tg), medial reticular nucleus (MRN) and parts of ventral tegmental nucleus (VTA). (**E**) Forebrain labelling appeared in zona incerta (ZI), lateral hypothalamus (LH), paraventricular hypothalamus (PVH), medial preoptic nucleus (MPO), stria terminalis (ST) and paraventricular thalamus (PVT). Scale bar: 500 μm (coronal sectional view) and 100 μm. (**F**) Distribution of fibers originating from RVLM_Dβh_ neurons across all areas labelled across the brain with color coding mapping to the spatial spread of the projections (5 mice).

With regards to labeling in brainstem areas away from the RVLM and caudal to the injection site, we observed that fibers surrounded the central canal within the dorsal motor nucleus of the vagus (DMV), nucleus tractus solitarius (NTS), and area postrema (AP). Labeling from the NTS extended rostroventrally into the medullary reticular formation, encompassing parvicellular (PCRt) and intermediate (IRt) subdivisions (**Figure 4C**). In brainstem areas rostral to the injection site, we observed dense labeling in the locus coeruleus (LC) (**Figure 4C**), with most fibers ipsilateral to the injection site, and only sparse contralateral projections, particularly to the LC.

With regards to labeling in the midbrain, we observed that rostral to the LC, fibers extended into the ventrolateral periaqueductal gray (PAG), the dorsal raphe nucleus (DR) (**Figure 4D**). Labelled fibers in the PCRt and IRt coursed rostrally through the subcoeruleus, converging in the laterodorsal parts of tegmental nucleus (Tg) and parabrachial nucleus (PBN) (**Figure 4D**). Additional labeling appeared in the medial reticular nucleus (MRN) and the lateral pontine nucleus (PN), which lie ventral to the adrenergic A7 group, appeared further rostrally in the retrorubral (RRF) and pararubral (PaR) fields, and appeared as sparse fibers in the ventral tegmental area (VTA) adjacent to TH-positive neurons (**Figure 4D**).

With regards to labeling in the forebrain, we observed dense projections in subthalamic and hypothalamic regions, including the dorsomedial (DMH), paraventricular (PVH), and lateral hypothalamus (LH), as well as the zona incerta (ZI) (**Figure 4E**). Fibers in LH spanned diversely rostrocaudally (**Extended Figure 3**). Additional fibers encircled the dorsal third ventricle (D3V) within the paraventricular thalamus (PVT) (**Figure 4E**). Fibers also passed through the rostral hypothalamic territories, such as medial (MnPO) and lateral preoptic areas (LPO), and extend further into the basal forebrain, particularly the stria terminalis (ST) and lateral septum (LS) (**Figure 4E**). Sparse labeling was also detected in the central medial thalamus and amygdala nuclei (**Extended Figure 4**).

The broad spatial spread of ascending projections from RVLM_Dβh_ neurons was quantified by counting the fiber coverage, in terms of (0.324 μm)^2^ pixels, across all labeled brain regions (**Figure 4F**). The borders of the regions were imposed with reference to the Paxinos stereotaxis atlas [55]. In cases where labeled axons crossed closely apposed nuclei along a continuous path, separating projections by individual regions would have required artificial truncation of the same axon tract. To preserve anatomical continuity and avoid over-interpretation, such nuclei were analyzed as grouped units: PCRt and IRt; PAG and DR; PBN and Tg; and ST and LS. Critically, although there are projections to areas that are pre-cortical, we failed to observe any direct projections from RVLM_Dβh_ neurons to cortex.

### RVLM_Dβh_ neurons project to the cortex via polysynaptic pathways

The determination of polysynaptic projections from RVLM_Dβh_ neurons to the cortex was a multistep process that starts with the use of viral transsynaptic tracer to detect candidate projections. A critical aspect of all viral transsynaptic tracers is that the timing and expression levels can depend on neuronal type and axonal length. Thus all candidates must be supplemented by auxiliary studies, followed by their verification through the inactivation of the candidate intermediate nuclei. We outline the overall logic for this process, and then provide details of the realization.

#### Logic

The first step involved cell-specific labeling with a transsynaptic anterograde label, *Cre*-dependent herpes simplex virus (HSV-H129-ΔTK-tdTomato) [56]. The strain HSV-H129 is predominantly an anterograde virus that will reliably jump across successive synapses [57]. We stereotaxically injected virus into the region of the RVLM of Dβh-Cre mice so that HSV-H129 was localized to RVLM_Dβh_ neurons.

The second step is to perform a multitude of parallel experiments on litter-mates, where the expression and transport of HSV-H129 is allowed to proceed for different time intervals (**Figure 5**). In practice, we chose survival times of 36-, 48-, 60-, and 72-hours (**Figure 5A-C**). We found that a period less than 36-hours was too short to observe sufficient labeling. A period of 72-hours led to labeling of cortical somata, a desired outcome, yet the survival probability of our mice declined markedly beyond 72-hours post-injection (**Extended Figure 5**). All animals were perfused after sacrifice and sectioned, the sections were counterstained, digitally imaged (**Figure 5D-F**), and labeled somata were manually counted (**Figure 6A**).

**Figure 5.**
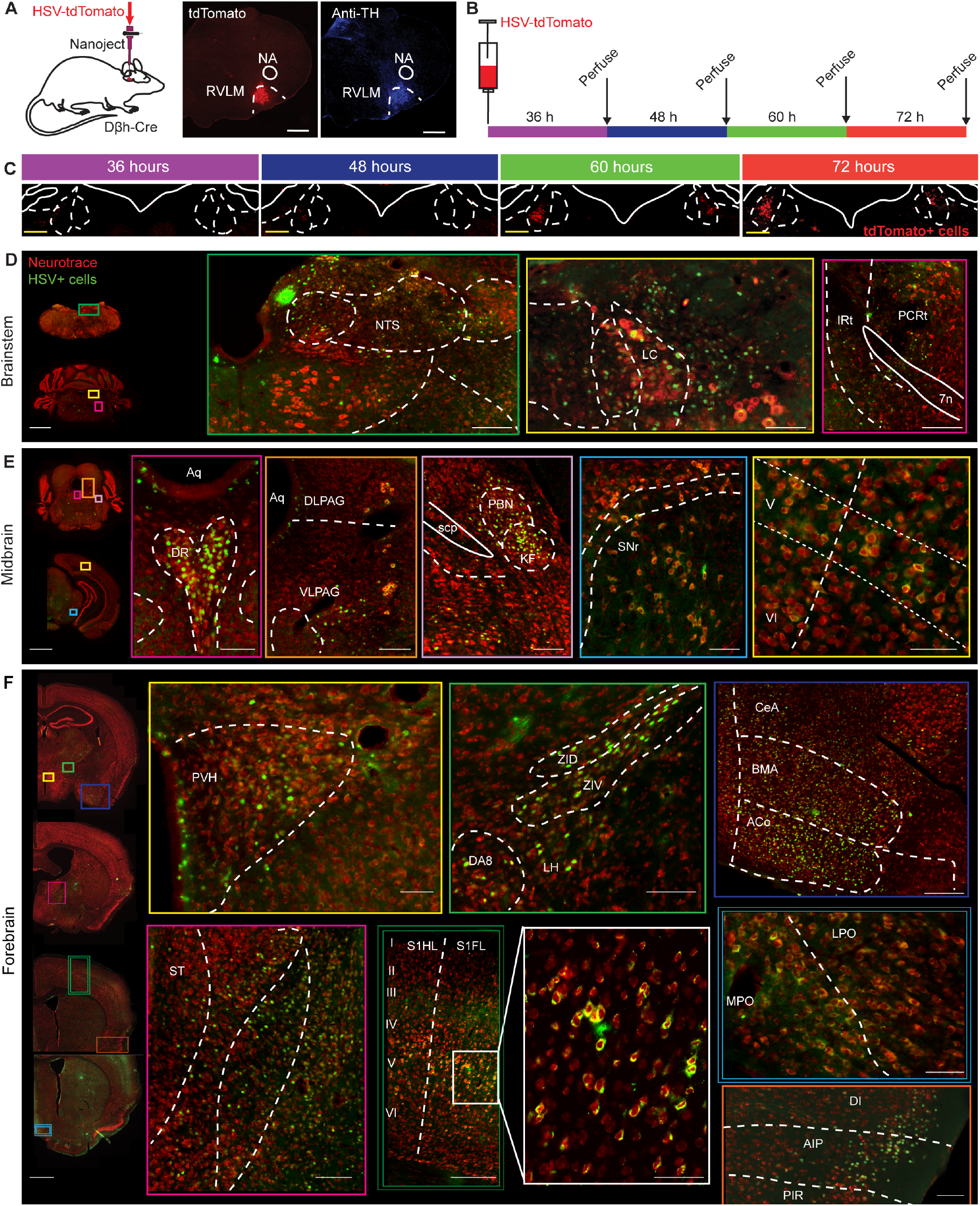
Polysynaptic circuits that involve RVLM_Dβh_ neurons.. (**A**) Schematic showing the injection site of HSV-H129-CAG-ΔTK-TT in a coronal brainstem section colabelled with tyrosine hydroxylase (TH) antibody. Scale bar: 500 μm. (**B**) Temporal labelling strategy illustrating timepoints of euthanasia at 36-, 48-, 60-, and 72-hour post injection. (**C**) Example images showing increase in cell labelling progressively with time in locus coeruleus (LC). Scale bar: 100 μm. (**D**) Brainstem labelling showing cells in NTS, PCRt, IRt, and LC. (**E**) Midbrain labelling showing cells in parabrachial nucleus (PBN), dorsal raphe (DR), venterolateral periaqueductal gray (VLPAG) and substantia nigra pars reticulata (SNr). (**F**) Forebrain labelling showing cells in dorsal and ventral zona incerta (ZID and ZIV), lateral hypothalamus (LH), paraventricular hypothalamus (PVH), lateral and medial parts of preoptic nucleus (LPO and MPO), parts of central amygdalar nucleus (CeA), basomedial amygdala (BMA) and anterior cortical amygdala (ACo), dorsal, agranular and piriform parts of insular cortices (DI, AIP and PIR), with large number of cells labelled in layer V of somatosensory cortex (S1Fl, S1DZ and S1BF) and sparsely in stria terminalis (ST). Scale bar: 500 μm (coronal sectional view) and 100 μm for zoomed in tiles.

**Figure 6.**
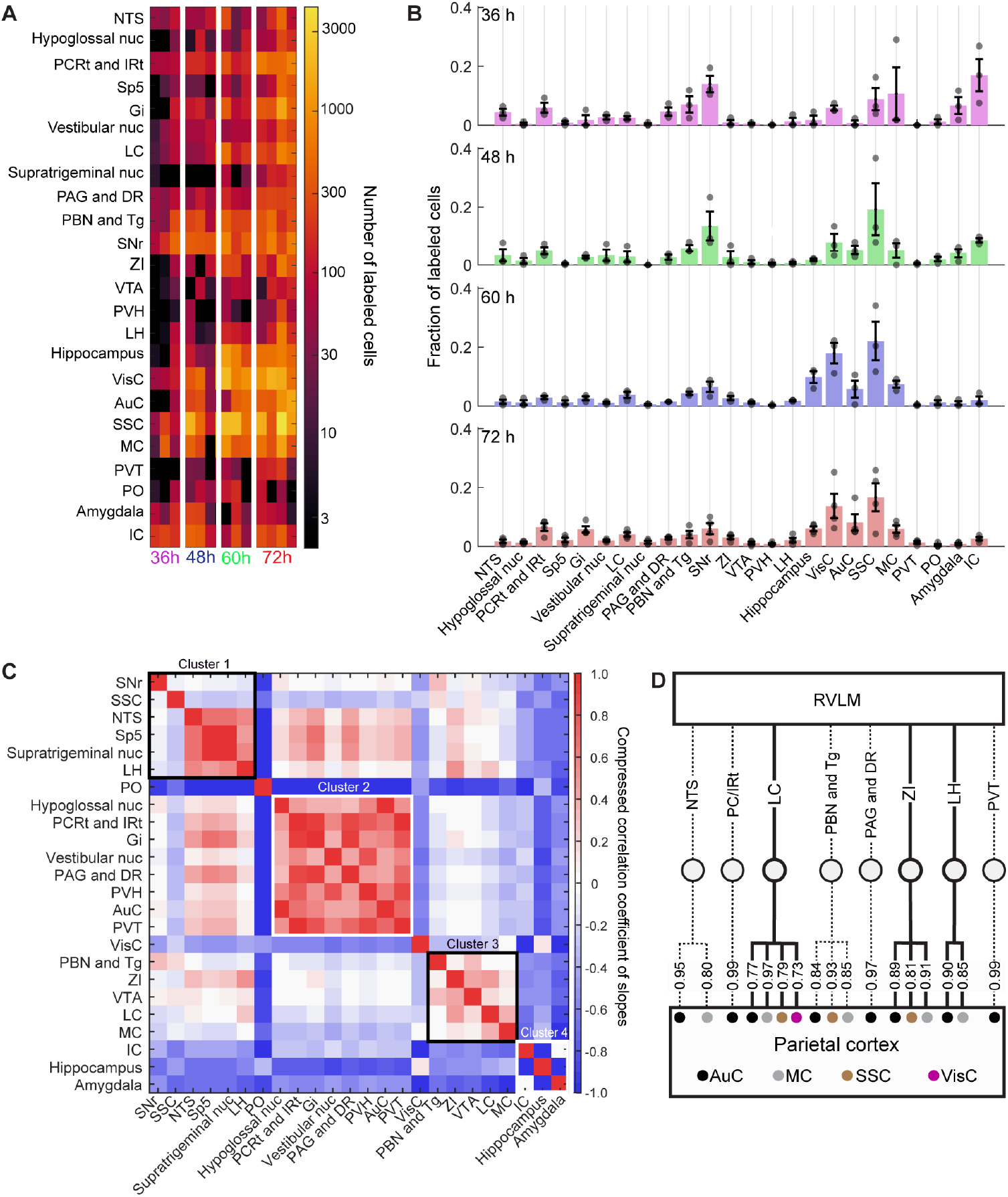
Disynaptic circuits from RVLM to parietal cortex. (**A**) Heatmap showing total number of cells labelled over time in all the areas when euthanized at 36 h, 48 h, 60 h, and 72 h. (**B**) Mean fraction of cells showing varying distribution of labelled cells at different time points (36 h: 3 mice, 48 h: 3 mice, 60 h: 3 mice and 72 h: 4 mice). (**C**) Cross correlation matrix (R) of the slope of cell labelling across different areas showing clusters of similar labelling profiles across areas. Matrix shown are Fisher Z-transform (tan^-1^R) and Z-score normalized for display. (**D**) Dendrogram of disynaptic circuits from RVLM to parietal cortex showing correlation coefficient values for each of the connected edge. Solid line shows the projections confirmed from the literature (**Extended Table 2**). Areas: nucleus solitaris (NTS), Hypoglossal nucleus (Hypogloss nuc), parvicellular and intermediate reticular nucleus (PCRt and IRt), Spinal trigeminal nucleus (Sp5), gigantiocellular nucleus (Gi), Vestibular nucleus (Vestibular nuc), Locus coeruleus (LC), Supratrigeminal nucleus (Supratrigeminal nuc), periaqueductal gray and dorsal raphe (PAG and DR), Parabrachial nucleus and Tegmental nucleus (PBN and Tg), substantia nigra (SNr), Zona incerta (ZI), ventral tegmental area (VTA), paraventricular hypothalamus (PVH), Lateral hypothalamus (LH), Hippocampus, Visual cortex (VisC), Auditory cortex (AuC), somatosensory cortex (SSC), motor cortex (MC), paraventricular thalamus (PVT), preoptic nucleus (PO), Amygdala, insular cortex (IC).

The third step is to employ a statistical metric to deduce likely polysynaptic path, or paths, defined as sequence of emergence of labeling of different nuclei. Our approach assumed that the expression of HSV-H129 continues to increase in the somata of a presynaptic brain nucleus as the virus jumps to a downstream postsynaptic nucleus. This assumption will hold when sequential nuclei make feedback as well as feed-forward projections and/or make recurrent connections within each nucleus. Phenomenologically, it appears to hold for all nuclei of interest (**Figure 6B**; **Extended Figure 7A**) and thus helps to narrow the list of candidate paths. The likely paths were assessed by calculating the slopes of somatic expression, assessed by the number of labeled cells between time points (**Extended Table 1**) and correlating the value of the slopes across all labeled brain nuclei (**Figure 6C**).

In the fourth step, we culled the candidates based on the results of prior anatomical analysis (**Extended Table 2**). (i) We selected the subset of candidate nuclei that were also significantly targeted by AAV labeling from RVLM_Dβh_ neurons (**Figure 4F**). (ii) We computed the total number of known of afferent sources and efferent targets of each candidate nucleus as a measure of its centrality [58]. Notably, hubs tend to have a large centrality (**Extended Figure 7D**,**E)**. (iii) We searched the literature (**Extended Table 2**) to confirm if candidate relay nuclei were reported to make connections to parietal cortex (**Figure 6D**). We pursued only nuclei that had at least one claim of a projection.

The results from the above four anatomical tracing and analysis steps yielded candidate pathways whose actual impact on CBF remained to be confirmed or dispelled. In the fifth step, we started with mice that expressed ChR in RVLM_*Dβ*h_ neurons and further made stereotaxic viral injections to express a chemogenetic inactivation agent [59, 60] in candidate brain nuclei that could serve as relays from RVLM to cortex. This permitted us to determine the impact of activating RVLM_*Dβ*h_ neurons on CBF before, during, and after inactivation of a potential relay neuron. Of course, there can be both multiple and overlapping pathways for RVLM_*Dβ*h_ neurons to dilate cortical arterioles.

#### Temporally controlled anterograde transsynaptic tracing

We observed labeling within the brainstem beyond expression in RVLM. Dense labeling was seen in LC, PCRt, IRt, and NTS, with sparser labeling in the gigantocellular nucleus, vestibular nucleus, hypoglossal nucleus, and spinal trigeminal nucleus (**Figure 5D**). This set of nuclei does not include the AP, which was labeled with anterograde AAV tracing (**Figure 4C**). In the midbrain, labeled neurons were found in the substantia nigra pars reticulata (SNr), PBN, DR, and ventrolateral PAG (**Figure 5E**). This labeling did not extend to the MRN, which was, however, labeled with anterograde AAV tracing (**Figure 4D**). In the forebrain, rostral labelling was prominent and included the insular (IC), layer V of somatosensory (SSC) and motor (MC) cortices, and sparser labeling in visual cortex (VisC) and auditory cortex (AuC) (**Extended Figure 6**). The broad distributions of transynaptically virus labelled cortical somata across the cortical mantle (**Extended Figure 6**) is consistent with the widespread activation of cortex by driving RVLM_Dβh_ neurons (**Figure 2F-J)**. Additional labeled cells were observed in the central amygdala nuclei (CeA) and in hypothalamic nuclei including PO, PVH, LH, and DMH, as well as ZI (**Figure 5F**). This set does not include the ST and the LS, which were labeled with anterograde AAV tracing (**Figure 4D**).

Four regions labeled by HSV-tracing did not overlap with AAV-defined primary projection sites. While the polysynaptic labeling did not include the projection of RVLM_Dβh_ neurons to the MRN nor AP, there is no reported evidence for MRN nor AP projections to cortex. Thus, it is unlikely that we missed identifying a potential disynaptic pathway that includes MRN or AP. Polysynaptic labeling also did not include the LS, which was weakly labeled with anterograde AAV tracing. The LS, along with the weakly labeled ST and PO, are part of the basal forebrain, so this could be part of a weak disynaptic projection to cortex.

#### Selection of potential pathways

Quantification of labeled cells across time points revealed a redistribution of labeling from subcortical to cortical regions between 36 and 72 hours, indicating progressive transsynaptic propagation to higher-order circuits (**Figure 6A,B**). We calculated the pathlength of all the labelled areas as the Euclidean distance from RVLM with reference coordinates estimated from the Paxinos atlas [55] and found that labeling expanded rostrocaudally from the RVLM (**Extended Figure 7B**). Next, we identified brain regions with similar growth rates of labeling (**Figure 6C**), as represented by a distance-based clustering of correlated nodes on a three-dimensional graph of RVLM_Dβh_ connectivity (**Extended Figure 7C**). This led to three putative clusters of nuclei, yielding 17 potential candidates for disynaptic pathways (**Figure 6C**).

We culled the candidates by excluded nuclei that were not labeled by both AVV (**Figure 4**) and HSV-H129 (**Figure 5**). This reduced the candidate pool to 12 nuclei. Next, we weighted each path by network centrality [58] (**Extended Figure7D; Extended Table 2**) (Methods). The dominant cluster of candidate nuclei had relatively large centrality yet led to a further cut that reduced the candidate pool to 10 nuclei (**Extended Figure 7D**,**E**). Lastly, we searched the literature and determined that only 3 of the 10 candidates were reported to make connections to parietal cortex (**Figure 6D**). Nuclei that were grouped in the anterograde analysis (**Figure 5**) were retained as groups during hierarchical clustering. This allowed the dendrogram to reflect pathway-level organization, while avoiding artificial subdivision of continuous axonal and transsynaptic routes (**Figure 6D**).

Our analysis yielded two anatomically and functionally distinct domains as disynaptic pathways. One domain is a brainstem pathway dominated by the LC and that also encompasses dorsal and ventral PAG and Tg. The LC is known to project to cortex [61]. The second domain is a midbrain/subthalamic pathway dominated by ZI and LH and that also encompasses PaR, SNr, SNc, and fields of Forel (F). Both LH and ZI are known to project to cortex [62-65] as intermediate nuclei.

### Chemogenetic inhibition of relay targets reveal a dominant disynaptic pathway

To functionally confirm the contribution of the potential pathways, we performed chemogenetic inhibition of the respective nuclei while measuring cortical vasodilation in response to optogenetic activation of RVLM_Dβh_ neurons (**Figures 3A** and **7A,B**). Two experimental strategies were employed: (1) Selectively labeling and inhibition of the individual pathways; and (2), concurrent inhibition of both pathways. In each mouse, we measured cortical vasodilation evoked by optogenetic stimulation of RVLM_Dβh_ neurons after administration of saline, as a control, and CNO (**Figure 7C-E**).

**Figure 7.**
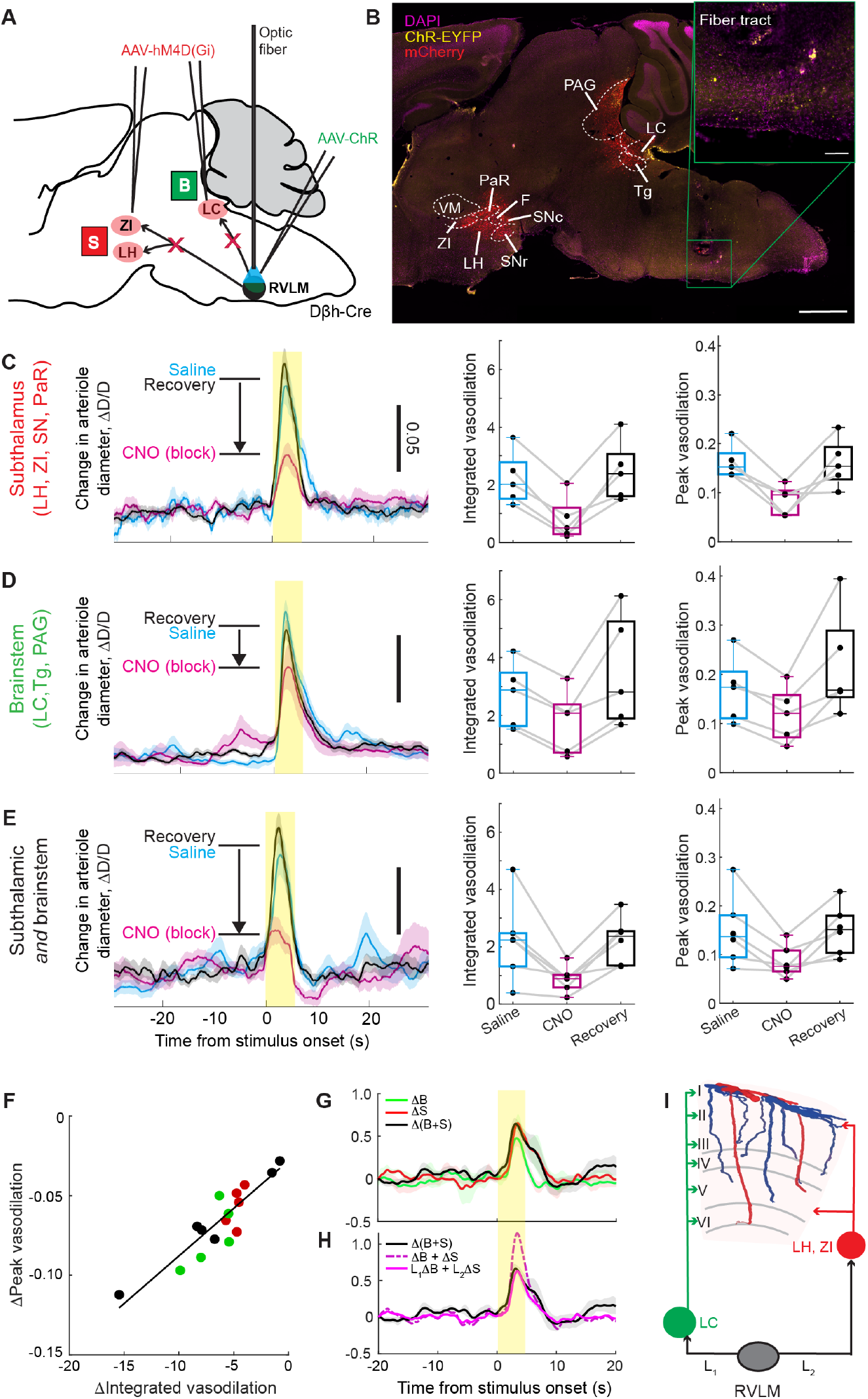
Chemogenetic inhibition of brainstem and subthalamic nuclei reduces RVLM_Dβh_-evoked cortical vasodilation. (**A**) Schematic showing injection sites of AAV8-hSyn-hM4D(Gi)-mCherry in locus coeruleus (LC), lateral hypothalamus (LH), and zona incerta (ZI) and injection of AAVDJ-EF1α-DIO-hChR2(H134R)-eYFP-WPRE-pA in *region* RVLM. (**B**) Sagittal section showing labelled cells at the injection sites and location of optic fiber. Note that subthalamic domain included labelled cells in pararubral area (PaR), fields of Forel (F), substantia nigra pars reticulata (SNr) and compacta (SNc) in addition to LH and ZI. Similarly, brainstem domain included periaqueductal gray (PAG), and tegmentum (Tg) in addition to LC. (**C-E**) Trial averaged RVLM_Dβh_-evoked vasodilation measured from cortical arteriole with two-photon microscopy while blocking subthalamic (C), brainstem (D) and both nuclei (E). Left panels illustrate example time series from single mouse showing the effect of clozapine N oxide (CNO) injection comparing to saline (control) and CNO washout (recovery) while stimulating RVLM_Dβh_ neurons. Middle and right panels compare mean responses to blocking the circuit. Peak and integrated vasodilatory responses when inhibiting subthalamic nuclei are: 0.163 and 2.20 (saline), 0.085 and 0.799 (CNO), 0.161 and 2.463 (recovery) or inhibiting brainstem nuclei are: 0.168 and 2.702 (saline), 0.118 and 1.754 (CNO), 0.220 and 3.507 (recovery) or both nuclei are: 0.149 and 2.226 (saline), 0.086 and 0.868 (CNO), 0.151 and 2.235 (recovery). Both integral and peak of vasodilatory responses are significantly reduced when blocking subthalamic nuclei (5 mice and 34 trials, integral: p = 0.008, Peak: p = 0.037) and weakly reduced when blocking brainstem (5 mice and 39 trials, Integral: p = 0.08, Peak: p = 0.017). Blocking both subthalamic and brainstem nuclei reduced the RVLM_Dβh_-evoked vasodilatory response by 70 % (6 mice and 40 trials, Integral: p = 0.036, Peak: p = 0.020). (**F**) Scatter plot comparing reduction in evoked vasodilatory responses from RVLM_Dβh_ neurons (peak vs Integral) between all blocking experiments. (**G**) Mean effect size quantified for blocking brainstem (*ΔB*) or subthalamic (*ΔS*) or both nuclei (*Δ(B+S)*). (**H**) Models predicting mean effect size of blocking both nuclei by linear addition (*ΔB + ΔS*), and weighted norm addition of blocking individual nuclei (*L*_*1*_*ΔB + L*_*2*_*ΔS*), showing nonlinear contribution of the two pathways to cortical vasodilation. (**I**) Circuit schema illustrating disynaptic pathways of CBF control from RVLM_Dβh_ neurons that relay through LC or LH and ZI.

Within 30 minutes of CNO injection, vasodilation in response to optogenetic activation of RVLM_Dβh_ neurons was reduced across all groups, whether the subthalamic site (**Figure 7C**), brainstem site (**Figure 7D**), or both sites (**Figure 7E**). In detail, we observed a large, significant decrease in vasodilation (5 mice, 34 trials; Integral: p = 0.008, Peak: p = 0.037, repeated measures ANOVA) induced by activation of RVLM_Dβh_ neurons when subthalamic nuclei were inhibited across mice. We observed a weaker but significant decrease in vasodilation (5 mice, 39 trials; Integral: p = 0.083, Peak: p = 0.017) when we inhibited brainstem nucleus LC. Further when we inhibited both nuclei, we observed a decrease in vasodilation like that for the subthalamic nuclei alone (6 mice, 40 trials; Integral p = 0.036, Peak: p = 0.020). In all cases, vasodilation returned in response to optogenetic activation of CNO washout to confirm reversibility (**Figure 7C-E**). The time for full recovery differed between groups: recovery for mice with single-sites inhibition occurred within 12 hours while mice with dual-site inhibition required up to 72 hours.

On average, the inhibition of solely subthalamic nuclei produced an ∼ 70 % reduction in vasodilation amplitude of pial arterioles. This implies a particularly strong contribution of the disynaptic pathway through LH and ZI (**Figure 7F**). As a control, we note that the diameter of pial arterioles was unchanged by CNO administration to block different potential relay pathways prior to optogenetic stimulation of RVLM_Dβh_ neurons (**Extended Figure 8**).

To evaluate whether the combined inhibition effect reflected additive or overlapping influences (**Figure 7G**), we compared the observed reduction in the dual-inhibition condition with predicted responses derived from linear and weighted distance norms based on the Euclidean path lengths of these regions from RVLM (**Figure 7H**). Linearity suggests that the LH/ZI-pathway is inclusive of the LC-pathway. The weighted distance-norm model, which normalizes the contribution of each pathway by a multiplicative factor proportional to the relative path length of each nucleus from the RVLM, provided a closer fit to the measured response. This suggests potential nonlinear contributions of the subcircuits from RVLM_Dβh_ neurons to cortex (**Figure 7I**). Regardless, we have identified a disynaptic pathway that accounts for 70 % of the dilatory action of increased activity of RVLM_Dβh_ neurons.

## DISCUSSION

We addressed the functional impact and anatomy of a brainstem pathway for the modulation of CBF. In awake mice, we establish variability in the activity of RVLM_Dβh_ neurons precedes an increase in CBF (**Figure 1J**), and that the amplitude of the variability is increased by hypoxia (**Figure 1D,E**), which is known to increases CBF To establish a causal link between the activity of RVLM_Dβh_ neurons and changes in CBF, we optogenetically drove RVLM_Dβh_ neurons and observed a concomitant dilation of individual cortical pial arterioles as well as a global increase in CBF (**Figures 2 and 3**). Systemic functions, including cardiac pumping and respiration, were unaffected upon activation of RVLM_Dβh_ neurons (**Figure 3**). These data support regulation of CBF by a medullary nucleus that provides an adaptive mechanism to rapidly increase cortical perfusion following an environmental drop in pO_2_.

Across our cohort, optogenetic stimulation of RVLM_Dβh_ neurons produced a 17 ± 9 percent increase in CBF, measured with LDF, relative to baseline (**Figure 3E**). How does this compare with the results for changes in blood flow in response to activation of other neurovascular processes? This increase in CBF measured here by optogenetic stimulation of RVLM_Dβh_ neurons is the same, within uncertainty, as the 20 ± 6 percent increase in CBF measured across the pial surface with LDF in response to vibrissa stimulation [66]. The increase in CBF measured here is potentially larger than the 9 ± 2 percent increase in response to vibrissa simulation found by particle tracking of red blood cells (RBCs) in pial arterioles, potentially larger than the 13 ± 3 and 6.5 ± 1.5 percent changes in CBF in response to forelimb and vibrissa stimulation, respectively [67], inferred from laser speckle contrast imaging, and similar to the 26 ± 4 percent change in CBF in response to optogenetic stimulation of cortical inhibitor neurons, also inferred from laser speckle contrast imaging. Lastly, change in CBF in response to optogenetic stimulation of RVLM_Dβh_ neurons matches the 19 ± 7 percent increase in the magnitude of flux of RBCs flowing through penetrating arterioles during ongoing vasomotion [39]. Thus RVLM can modulate CBF to the same extent as other neurovascular processes.

The widespread activation of cortex by driving RVLM_Dβh_ neurons **Figure 2F-J** is supported by complementary anatomical findings that demonstrate broad distributions of transynaptically virus labelled cortical somata across the cortical mantle (**Extended Figure 6**). Labeling is seen as early as 36 hours after infection and peaks at 72 hours (**Figure 6F**). The results of the anatomical projection studies (**Figures 4 to 6**), assessed and augmented by chemogenetic inactivation of putative intermediate nuclei (**Figure 7**), provides direct evidence of disynaptic pathways from RVLM_Dβh_ neurons to neurons in cortex that regulate CBF. One such disynaptic pathway, involving the LH and ZI, accounts for 70 % of the regulation of CBF vis activation of RVLM_Dβh_ neurons. Notably, LH and ZI, along with the other major intermediate, the LC, are known integrative hubs [68-70] with an exceptionally large number of inputs and outputs (**Extended Figure 7D**,**E; Extended Table 2**). The involvement of LC is particularly notable given its role in attention [71, 72] and arousal [73] and the role of noradrenaline in shaping changes in cortical blood flow [15, 16, 74].

### A brainstem circuit that drives cortical blood flow

Reis and Golanov proposed a trisynaptic pathway with two intermediary brain nuclei that linked the RVLM to the control of CBF; the medullary cerebrovascular area (MCVA), located caudal to RVLM and ventral to nucleus ambiguous, and the subthalamic vasodilator area (SVA), encompassing the ZI, fields of Forel, and prerubral regions [21]. Their evidence was derived from lesion and stimulation experiments in anesthetized rats. First, electrical stimulation of the RVLM increased cortical δ-band oscillations in electroencephalogram (EEG) activity, suggesting involvement of thalamocortical circuits known to generate δ-rhythms [75]. Second, the evoked δ-band EEG activity after stimulation of the SVA occurred approximately 9 ms earlier than stimulation of the MCVA, suggesting the SVA is downstream of the MCVA. Third, lesions of the SVA halved hypoxia-induced vasodilation that was driven by RVLM activation [21]. Our findings revealed a simpler, disynaptic projection from the RVLM region (**Figure 7I**).

We do not observe an increase in δ-band power secondary to stimulation of RVLM_Dβh_ neurons (**Figure 3C**), as reported previously [76]. This difference could reflect our use of awake mice as opposed to anesthetized rats. It could also reflect our specific activation of adrenergic neurons (**Figure 4A,B**) in contrast to electrical stimulation; the latter procedure drives both orthodromic and antidromic potentials across mixed neuronal populations.

### Local and global control of CBF regulation

We found that increases in CBF that follow stimulation of RVLM_Dβh_ neurons precedes changes in the ECoG (**Figure 3D**). This implies that RVLM-dependent control of CBF is only weakly coupled to local neuronal activity in cortex (**Figure 3E**), whereas strong coupling through sensory stimulation leads to a local change in neuronal activity that precedes a change in flow by 2 s [77]. How does this occur? Fast components of vasodilation are driven by activity-dependent K^+^ efflux from potentially all neurons [8], as well as by interneurons that express neuronal nitric oxide synthase [10, 11]. It is thus possible that the dominant disynaptic pathways project to only selected neurons, although direct projections to arteriolar smooth muscle are also a possibility [13].

When systemic oxygen levels fall, sustained perfusion of the brain likely requires mechanisms beyond those active during normal homeostasis. This could be accomplished by changes in slow neurovascular responses, including the release of vasoactive intestinal protein, neuropeptide Y, somatostatin, and prostaglandins E, all of which prolong the hemodynamic response [5, 9, 11, 78, 79]. Further, long-range neuromodulation by the cholinergic basal forebrain, serotonergic raphe, noradrenergic LC, and dopaminergic VTA can sustain CBF by altering local circuit excitability and, in some cases, by acting directly on the vasculature [13, 15, 80-82]. Our data support a brainstem-initiated mechanism that dilates cortical arterioles independent of changes in cortical field activity, e.g., this could be mediated by release of orexin/hypocretin from LH and noradrenaline from LC.

A final remark is that brief optogenetic activation of RVLM_Dβh_ neurons increased CBF without affecting respiration or heart rate (**Figure 3**). This points to an intracerebral branch of the RVLM circuit potentially distinct from its descending sympathetic pathway.

### RVLM activity and hypoxia

While brainstem neurons are generally resilient to mild hypoxic events, prolonged or chronic hypoxia can lead to maladaptive cardiovascular outcomes. Notably, lesion studies show that ablation of RVLM neurons diminishes cerebrovascular responsiveness to hypoxia [83], underscoring their involvement in homeostatic vascular control.

Following reoxygenation, CBF rapidly returned to baseline; however, the variability in intracellular calcium activity of RVLM_Dβh_ neurons remained elevated for the duration of the recovery period (**Figure 1F,I**). This dissociation in activity implies that neural activity in key autonomic centers recovers slowly compared to systemic physiological effects. This is consistent with the phenomenon of “hypoxia hangover”, where peripheral measures stabilize within minutes, yet neural and cognitive functions may remain perturbed for hours [84, 85]. Hypoxia represents a bimodal physiological challenge [86]. Mild, intermittent hypoxia, such as occurs during exercise or altitude adaptation, can have beneficial cardiovascular and cognitive effects [87-89]. In contrast, chronic or severe hypoxia is detrimental to brain function [90-92]. Recent studies support hypoxia as a therapeutic method [93] that can provide neuroprotection in rodent models of Parkinson’s disease [94] and Alzheimer’s disease [95]. It is plausible that the threshold for tolerance to hypoxia, in terms of neuronal death, depends on the capacity of the RVLM_Dβh_ neurons and their downstream intracerebral circuits to compensate for fluctuations in pO_2_. In this sense, the greater RVLM may act to dynamically adjust cerebral perfusion on the slow time scales of hundreds of seconds, or longer.

### Epilogue

Our findings highlight the need to consider non-local effects in the regulation of cortical blood flow. Firstly, we emphasize the role of RVLM_Dβh_ neurons as an interoceptive interface between the peripheral and central circuits that adjust cortical perfusion in response to gas challenges. Secondly, we emphasize caution in the interpretation of functional imaging studies based on blood level oxygen dependent (BOLD) signals [96, 97]. It is typically assumed that changes in blood oxygenation are either of local origin or uniformly global [98]. Yet, activation of RVLM_Dβh_ neurons leads to changes in flow across cortex that maintain a spatial pattern independent of sensory stimulation or ideation (**Figure 2F-J**).

## Online methods

### Animals

All experimental procedures were approved by the Institutional Animal Care and Use Committee at the University of California, San Diego. Adult *Dβh Cre* mice [*Tg(Dβh-Cre)KH212Gsat*] were obtained from the Mutant Mouse Resource and Research Center (MMRRC) and maintained on a C57BL/6J background. Both male and female mice were used for experiments. The strain used for widefield imaging was a transgenic line; B6.Cg-Tg(Acta2-GCaMP8.1/mVermilion)B34-4Mik/J expressing GCaMP8.1 in smooth muscle-cells (JAX 032887).

### Viral vectors

AAV8-EF1α-DIO-mCherry (1.53 × 10^13^ GC ml^−1^) used for anterograde tracing and AAVDJ-EF1α-DIO-hChR2(H134R)-eYFP-WPRE-pA (1.22 × 10^12^ GC ml^−1^) for optogenetic activation of RVLM_Dβh_ neurons were obtained from the Sanford Viral Vector Core (Salk Institute, USA). HSV-H129-CAG-ΔTK-TT was obtained from the Center for Neurogenetic and Neurocircuitry Viral Vector Core (University of North Carolina, Chapel Hill, USA). Red shifted AAV8-EF1α-DIO-ChRmine-eYFP-WPRE was used for stimulation of the RVLM_Dβh_ neurons via the auditory canal coupled with widefield imaging and was obtained from Stanford viral core. Addgene-sourced vectors included AAV9-CAG-Flex-jGCaMP8s-WPRE-SV40 (1 × 10^13^ GC ml^−1^; plasmid #162380-AAV9) for calcium imaging, AAV8-hSyn-hM4D(Gi)-mCherry (7 × 10^12^ GC ml^−1^; plasmid #50475-AAV8) for chemogenetic inhibition, and AAV2-EF1α-DIO-eYFP (3 × 10^12^ GC ml^−1^; plasmid #27056-AAV2) as a control virus for ChR responses.

### Stereotaxic surgery

All surgical procedures were performed in 8-12-week-old mice under isoflurane anesthesia at 4 % (v/v) for induction, 1-2 % (v/v) for maintenance from a precision vaporizer (HME® vaporizer). Depth of anesthesia was monitored continuously via reflex testing and respiration rate. Body temperature was maintained at 37 °C using a feedback-controlled heating pad (FHC®, DC Temperature Controller). Mice were placed in a stereotaxic frame (Kopf Instruments) equipped with a micromanipulator (Sutter MP-285). Bregma and lambda were identified and leveled in both the mediolateral and anteroposterior planes according to the Paxinos mouse brain atlas [55].

#### Brainstem targeting

Viral injections targeting region RVLM were performed using a Nanonet III injector (Drummond Scientific, USA) fitted with pulled glass micropipettes (tip length ≈6 mm). Injections were delivered at a rate of 10 nl min^−1^, and the pipette was left in place for 40 minutes to allow diffusion before withdrawal. Stereotaxic coordinates for region RVLM were optimized using electrical stimulation to confirm CBF responses (**Extended Figure 9A-C**) and corresponded to bregma –6.75 mm, mediolateral 1.25 mm, and dorsoventral 4.9 mm from the brain surface. To identify the precise coordinates of the RVLM region for activation, we first performed electrical stimulation using bipolar electrodes (FHC) at 60 μA for 5 s at multiple sites along the brainstem under anesthesia and monitored cortical perfusion using LDF. Maximal increases in CBF were observed over the parietal cortex (**Extended Figure 9**), consistent with coordinates, Bregma-6.75 mm, mediolateral 1.25 mm and dorsoventral 4.9 mm from the brain surface.

#### Thinned-skull window and optic fiber implantation

Mice underwent a second surgery 4 to 5 weeks post viral injection, for transcranial window preparation and optic fiber implantation. The scalp was removed under aseptic conditions, brain surface levelled to align bregma and lambda as described earlier, followed by <1 mm craniotomy using a 250 μm diameter burr in a low-vibration drill (Osada EXL-M40) at the coordinates of injection. Custom-made optic fibers (0.48 NA, 6 mm length, cut and polished at a 90° angle; Doric®) were then inserted into the site of injection and fixated using cyanoacrylate glue (Loctite 401). The periosteum over the parietal and occipital bones was gently cleared. Skull sutures were reinforced with low-viscosity cyanoacrylate glue (Loctite 4104). For all experiments with fiber implantation, a ∼4 mm diameter region centered over the primary vibrissa cortex (bregma –1.5 mm, mediolateral 2 mm) was thinned using a 250 μm burr (Osada EXL-M40) to form a chronic transcranial window according to the methods described previously. For the cortical imaging of smooth muscle-cell calcium a region spanning 10 mm x 10 mm region of the cortex was thinned [39]. The thinned region was dried, coated with cyanoacrylate (Loctite 401), and sealed with a No. 0 glass coverslip. A custom metal head bar was affixed to the interparietal bone using dental cement (Metabond) to stabilize the preparation for imaging.

### Fiber photometry

For calcium imaging, Dβh-Cre mice received stereotaxic injections of AAV9-CAG-Flex-jGCaMP8s-WPRE into the RVLM. After 4 weeks of viral expression, an optic fiber was implanted at the injection site, and mice were allowed to recover for 5 days. Following recovery, animals were habituated to the combined fiber photometry and LDF setup for 4 consecutive days before the day of recording.

The fiber photometry system consisted of a custom-built hybrid configuration using an IIFMC4 fluorescence cube (Doric®) and was coupled to an Arduino-controlled LED driver to multiplex 415 nm (isosbestic) and 488 nm (GCaMP) excitation LEDs at 200 Hz. Emission (500-540 nm) was collected through a low-autofluorescence patch cord, and the cube gain was set to 10-times amplification. The combined excitation signals were digitized and used post hoc to demultiplex the emission signal using custom MATLAB scripts that extracted the isosbestic (415 nm) and calcium-dependent (488 nm) components for further analysis (**Eqns 1**,**2**).

### Hypoxia experiments

A custom hypoxia chamber was fabricated using a laser-cut acrylic enclosure fitted with inlets for gas delivery of room air and 100 % (v/v) N_2_ and outlet for vacuum. Oxygen levels within the chamber were modulated manually and measured using I^2^C oxygen sensor and logged to Python interfacing via an Arduino. Carbon dioxide concentration was measured with a K30 sensor (GasLab) and read directly onto the computer.

During each experiment, [Ca^2+^] from RVLM_Dβh_ neurons along with LDF over the parietal cortex were recorded simultaneously. Each session consisted of 50 minutes of baseline recording under normoxia, followed by 30 minutes of hypoxia, and 50 minutes of recovery. The transition to hypoxia was defined by oxygen concentration falling below 16 % (v/v), as determined by the oxygen sensor.

### Laser Doppler flowmetry

CBF was monitored using a MoorLab LDF system (Moor Instruments, Millwey, Axminster, Devon, UK) operating at a wavelength of 780 nm and low pass filtered at 15 kHz, yielding a maximum measurable velocity of λ_0_f_cut/2 = 6 mm/s. A fiber-optic MP1-V2 probe was positioned above the thinned-skull window using a custom adapter. The LDF *flux* signal, representing the product of average red blood cell velocity and backscattered light intensity, was acquired at 1 kHz using Labchart software (ADInstruments®).

### Optogenetic stimulation of RVLM_Dβh_ neurons

Following surgery, mice were allowed to recover for 5 to 7 days and were then habituated to head fixation for 4 consecutive days before imaging. To optogenetically activate RVLM_Dβh_ neurons, 488-nm laser light was delivered through an implanted optic fiber. Multiple stimulation frequencies: 10, 30, 50 Hz and a continuous pulse of 5 s (**Figure 3C**) and power levels were tested to establish an effective stimulation of RVLM_Dβh_ neurons in vivo (**Extended Figure 10**). We chose 5 s of continuous square-wave pulse (8–12 mW at the fiber tip) for our two-photon microscopy experiments, as it reliably evoked cortical arteriolar dilation (**Figure 2**).

### Electrophysiology

Two 50 μm diameter tungsten Teflon coated wires (AM Systems, no. 794-623) were implanted during the second surgery post 4 to 5 weeks of viral injection. The electrodes spanned the thinned skull window and were inserted to touch the cortical surface. A third electrode was inserted deep in the cerebellum and served as a reference. All electrodes were stripped of 1mm of insulation before implantation to expose the conducting end. The ECoG signal was amplified (World Precision Instrument, DAM 80), filtered between 0.1 Hz and 10 kHz, digitized, and stored for post hoc analysis.

### Physiology measurements (breathing and heart rate)

Breathing rate was monitored using a far infrared thermal imaging camera (FLIR Systems®) positioned to capture temperature fluctuations at the nostrils of mice while stimulating RVLM_Dβh_ neurons. Nostril motion was first corrected using DeepLabCut®, and the average temperature across both nostrils was extracted using custom MATLAB scripts. Breathing frequency was computed from the averaged nostril temperature signal by high-pass filtering (cutoff = 1 Hz), applying a zero-phase gaussian filter, and detecting individual breathing peaks. Interbreath intervals (IBIs), representing the time between consecutive breaths (**Eqn 4**), were calculated, and averaged across prestimulus (Pre), stimulation (Stim), and post-stimulation (Post) epochs. Spectrograms were generated using the Chronux toolbox [99] to visualize breathing frequency over time.

Heart rate was calculated from recorded electrocardiogram (ECG) measured with electrodes (Model 2560, 3M Red Dot Monitoring Electrodes) attached to a custom fabric hammock on which mice were head fixed. Animals were habituated to the hammock for approximately 10 days before recordings to minimize motion artifacts. ECG signals were recorded across the forepaws, preamplified, and digitized while stimulating RVLM_Dβh_ neurons. Data were processed, and R-R interval was calculated (**Eqn 5**) using custom MATLAB scripts.

### Two-photon microscopy of cortical vessels

Cortical vessels were imaged transcranially through the thinned skull over the parietal cortex in awake, head-fixed mice, as described previously [100] Imaging was performed using an adaptive optics two-photon microscope to correct for the aberration. A 4-mm working-distance water-immersion objective (Olympus, XLPLN25XSVMP2; 25×, 1.0 NA) was used for standard imaging, whereas a 7.77-mm working-distance air objective Thorlabs, TL10X-2P; 10X, 0.5 NA) was used when needed to avoid potential collisions with the implanted optic fiber. To visualize the vasculature, mice received a retro-orbital injection of Cy5.5-conjugated 2000-kDa dextran 20 minutes before imaging. Cy5.5 fluorescence was excited using a 1250-nm wavelength at 20 to 40 mW, and the emitted fluorescence was bandpass-filtered (Semrock, FF01-708/75-25) before being collected by a silicon photomultiplier (Hamamatsu, C13366-3050GA). Arterioles and veins were distinguished based on the direction of flow and the presence of vasomotion. To prevent optogenetic stimulation artifacts in the two-photon data, the stimulation was gated only during the corner-scanning periods, which lie outside the temporal window used for fluorescence acquisition (∼30 % of the imaging cycle).

### Widefield microscopy

Awake head-fixed mice expressing the genetically encoded calcium indicator GCaMP8.1 in smooth muscle-cells were imaged using a Zeiss SteREO Discovery.V8 microscope with ThorLabs M470L3 excitation LED and Zeiss 38 HE filter-set (470/40 nm excitation bandpass filter, 495 nm dichroic, and 525/50 nm emission bandpass filter). Images were collected at ∼5 Hz with a Prime95B 1200 x 1200 sCMOS camera (Teledyne Photometrics). Stimulation of RVLM_Dβh_ neurons was performed using the red shifted opsin ChRmine [40] and placing a 630 nm LED (Mouser 941-XBDRED0701) with a total power of 100 mW at the opening of the external auditory canal, thus illuminating the brainstem at a distance of 8 to 10 mm from the midline through intact tissue and bone as described [101]. Ultra-slow stimulation was delivered at 0.14 Hz or 0.2 Hz for 500 s; pulses were 50 Hz at 50 percent duty cycle and 2 s duration. Arterioles were masked and segmented as described [39], and summary plots were made using the average GCaMP8.1 signal from all arterioles in the field of view. The GCaMP8.1 time and amplitude corresponding to peak dilation (negative ΔF/F) were calculated from each arteriole segment’s stimulus-triggered average signal and plotted across space.

### Anterograde monosynaptic and polysynaptic tracing

For anterograde tracing, Dβh-Cre mice received stereotaxic injections of AAV8-EF1α-DIO-mCherry into region RVLM. Brains were collected 4 to 5 weeks post-injection, perfused, and post-fixed in 4 % (w/v) paraformaldehyde. Coronal sections were prepared, mounted, and imaged sequentially using a slide scanner. Fluorescently labeled axonal projections were quantified across brain regions using custom MATLAB scripts to segment mCherry-labeled fibers from every third section of the brain followed by pixel quantification for each of the labelled areas. Brain areas were identified using Allen and Paxinos brain atlases [55].

For polysynaptic circuit tracing, mice were injected with an anterograde herpes simplex virus (HSV-H129-CAG-ΔTK-TT) [56] in region RVLM. Animals were perfused at survival times; 36, 48, 60, and 72 h post-injection to capture temporal stages of viral spread. Brains were sectioned, stained, mounted, and imaged, to assess the labelling across the brain. Cell counting was performed manually (**Eqn 6-8**) and analyzed in MATLAB using custom-written scripts.

### Chemogenetic inhibition of the circuit

For chemogenetic inhibition of intermediary relay nuclei, AAV8-hSyn-hM4D(Gi)-mCherry was stereotaxically injected into the LH (bregma –2.0 mm, mediolateral +1.0 mm, dorsoventral –4.5 mm), ZI (bregma –2.0 mm, mediolateral +1.0 mm, dorsoventral – 3.8 mm) or LC (bregma –5.4 mm, mediolateral +0.8 mm, dorsoventral –3.5 mm) or both. In the same animals, *Cre*-dependent AAVDJ-EF1α-DIO-hChR2(H134R)-EYFP was injected into the region of RVLM. Injection volumes (70 to 100 nl) were optimized to encompass the target nuclei and adjacent regions.

Four weeks after viral expression, mice underwent a second surgery for implantation of an optic fiber above the injection site as described earlier, followed by skull thinning over the parietal cortex for two-photon imaging. Following 5 days of postoperative recovery, animals were habituated to the imaging setup for 4 consecutive days. Control measurements were obtained after *i*.*p*., saline injection, during which vasodilation induced by photostimulation of RVLM_Dβh_ neurons was imaged, and the same vessel locations were marked for subsequent sessions.

Clozapine-N-oxide (CNO; 10 mg kg^−1^, *i*.*p*.) was then administered, and imaging was repeated 20-minute post-injection to assess the effects of chemogenetic inhibition on cerebral vasodilation. Recovery of vascular responses was evaluated 12 to 72 h after CNO administration. Recovery times varied depending on the targeted nuclei, with approximate recovery at 12 h for subthalamic inhibition, 24 h for brainstem inhibition, and 72 h when both nuclei were simultaneously inhibited.

### Pharmacology

Sodium nitroprusside (Sigma-Aldrich) was administered via retro-orbital injection following a 15-minutes baseline recording. After a 10-minutes interval to allow for drug onset, CBF and RVLM_Dβh_ [Ca^2+^] were recorded for an additional 30 minutes.

### Perfusion and histological preparation

At the conclusion of experiments, mice were anesthetized and transcardially perfused with phosphate-buffered saline (PBS) followed by 4 % (w/v) paraformaldehyde in PBS. Brains were dissected, post-fixed for 30 to 60 minutes at room temperature, stored overnight at 4°C in PFA, and subsequently transferred to 30 % (w/v) sucrose in PBS for 12–16 h. Serial 30-μm coronal sections were cut on a freezing microtome (MICROM® HM 400).

Sections were rinsed in PBS and processed for immunofluorescence or counterstaining to verify viral expression, injection sites, and fiber placements. Immunostaining for TH (1:1000, Millipore, AB152) was used to identify adrenergic neurons, followed by counterstaining with NeuroTrace 435/455 Blue fluorescent Nissl stain (1:200, Thermo Fisher Scientific). Sections were wet-mounted, and cover slipped using Fluoromount-G (SouthernBiotech®). In some cases, DAPI Fluoromount (SouthernBiotech®) was used in place of NeuroTrace.

### Data analysis

#### Calcium signal processing

Fiber photometry signals were first demultiplexed into the control (415 nm) and calcium-dependent (488 nm) channels. To correct for bleaching and motion artifacts, the 415 nm control signal was linearly fitted to the 488 nm calcium-dependent signal. The normalized fluorescence change (ΔF/F) was then computed as:

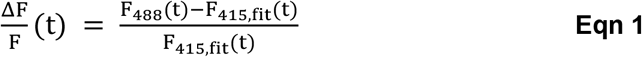

where F_488_ represents the demultiplexed 488 nm evoked fluorescent GCaMP signal and F_415_,_fit_ is the fitted isosbestic 415 nm evoked signal. Mean of baseline ΔF/F was subtracted from as shown in **Eqn 2**.

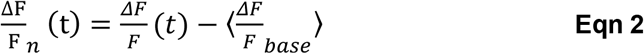

#### LDF quantification

LDF signal was used to calculate fractional change from baseline to compare across mice and is as follows:

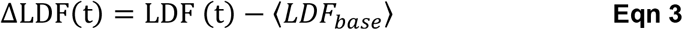

where LDF(t) represents the instantaneous laser-Doppler signal at time *t*, and ⟨*LDF*_*base*_⟩ denotes the mean LDF value during the baseline period prior to optogenetic stimulation of RVLM_Dβh_ neurons or hypoxia.

For calculating the cross correlation between ΔF/F and LDF, the modulation of oxygen was removed using a median filter fitted to the oxygen sensor signal followed by calculation of cross correlation for three time periods defined as 40 minutes before hypoxia (baseline), 30 minutes after oxygen levels drop below 19 % (hypoxia) and 40 minutes after oxygen levels resume to 21 % (v/v) (recovery).

#### ECoG signal analysis

Electrocorticogram (ECoG) signals were band-pass filtered between 1 and 100 Hz to remove slow drifts and high-frequency noise while preserving physiologically relevant activity. To examine spectral content, power spectral density (PSD) was estimated using the multitaper method from the Chronux toolbox [99]. Time to peak were calculated to assess the temporal order of onset of changes in CBF and ECoG under stimulation. Gamma band power was calculated as the average spectral power in the range of 30-60 Hz.

#### Breathing rate calculation

Breathing dynamics were extracted from thermal recordings obtained using a far infrared (FLIR®) infrared camera directed at the animal’s nostrils. The temperature trace was first smoothed using a low-pass filter (cutoff < 10 Hz) to reduce high-frequency noise and then breathing peaks were automatically detected from the temperature oscillations using a peak-finding algorithm in MATLAB. The IBI was defined as the time difference between consecutive breathing peaks:

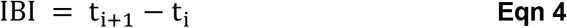

where t_i+1_ and t_i_ are the timestamps of successive detected peaks. From these values, mean IBI, breathing frequency. Peak amplitude was calculated from the detected peaks were computed across trials to quantify respiration patterns before (Pre), during (Stim), and after (Post) stimulation.

#### ECG analysis

Electrocardiogram (ECG) signals were band-pass filtered between 3 and 25 Hz to remove baseline drift and high-frequency noise while preserving the Q-R-S complex. R-wave peaks were automatically detected using MATLAB, with empirically defined thresholds for peak prominence and minimum interpeak distance based on the physiological range of mouse heart rates. The upper limit for detection was set to 800 beats per minute (bpm) and the lower limit to 400 bpm, ensuring that only physiologically plausible R–R intervals were included in the analysis.

The R–R interval was calculated as the time difference between consecutive R peaks:

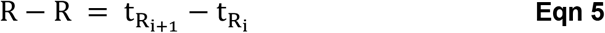

where 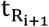 and 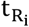 represent the timestamps of successive R-wave maxima. Instantaneous heart rate was computed as the R–R interval over 10 s overlapping windows, and mean R-R was averaged across trials and experimental conditions.

#### Diameter quantification

Cortical vessel diameters were quantified from two-photon imaging data. Each vessel was first segmented using intensity thresholding to enhance contrast and generate clear vessel boundaries. The vessel diameter was then calculated as the full width at half maximum of this averaged profile using custom MATLAB scripts.

#### Cell quantification for HSV cell labelling

For each animal, cell counts were obtained from manually annotated regions defined using area boundaries from the Allen Brain Atlas and Paxinos [55]. The total number of labeled cells was quantified for each brain area and time point (36, 48, 60, and 72 h post-injection). To normalize across animals, fractional labeling for each region and time point was calculated as:

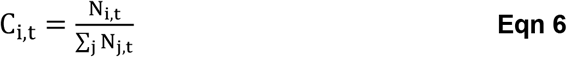

where N_i,t_ represents the number of labeled cells in region *i* at time *t*, and ∑_j_ N_j,t_ is the total number of labeled cells across all regions in that mouse at the same time point. To quantify temporal propagation of labeling, mean cell-labeling slopes, 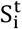, were computed for each region as the rate of change in cell number between consecutive time points, divided by the 12-hour interval:

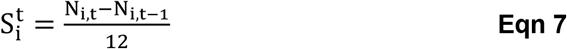

We next calculated a cross correlation between different areas and plotted the Pearson’s correlation coefficient, r_i,j_ between regions to find areas that show similar slope.

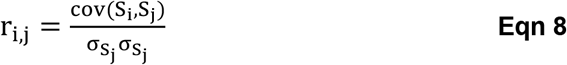

The resulting Pearson correlation matrix, R was converted to a distance matrix (1 − R) which was then used to cluster regions with similar temporal labeling profiles (**Figure 6C**). Data were then reordered by clusters to visualize structure within the correlation matrix. Pairwise regional correlation matrices were Fisher *z*-transformed using:

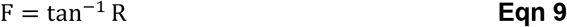

and z-scored for visualization to linearize correlation variance and enhance contrast across the matrix using:

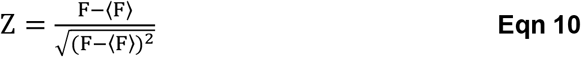

Regions exhibiting strong correlations (r > 0.7) were first identified as potentially connected areas. These regions were then subjected to a multi-step selection process to nominate candidates projecting to parietal cortex and hence targets for chemogenetic targeting:

1. **Anatomical validation:** From the set of highly correlated regions, we identified those labelled in AAV anterograde tracing experiments to confirm their status as primary projection targets.
2. **Network centrality analysis:** For this anatomically validated subset, we quantified network degree as the sum of known afferent and efferent connections, compiled from published anatomical literature (**Extended Table 2**). Regions were clustered based on this centrality measure, and those exhibiting relatively high centrality were selected using a threshold (**Extended Figure 7**).
3. **Cortical projection specificity:** Finally, from the high-centrality candidates, we identified regions that project to the parietal cortex, the site of CBF measurements, and selected these regions for chemogenetic inhibition.

#### Effect size quantification for chemogenetic experiments

For each animal, vessel diameter changes ΔD/D were extracted as time traces for the saline, CNO, and recovery conditions. To quantify the effective change in vasodilation following chemogenetic inhibition, the normalized difference between saline- and CNO-evoked responses was computed as:

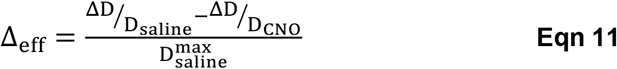

where 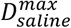 represents the maximum dilation measured in the saline condition. This metric was calculated for chemogenetic blocking of brainstem, i.e., ΔB, subthalamic, i.e., ΔS, and both nuclei, i.e., Δ(B + S). The resulting values were compared across groups to assess the relative contribution of each circuit cluster to RVLM_Dβh_ -evoked vasodilation.

To evaluate whether the combined inhibition effect could be derived from data of inhibition the individual nuclei, following two predictive models were tested:

1. **Linear model**:

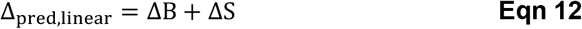
2. **Weighted norm model**{

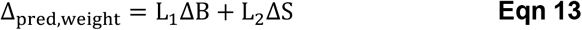

where scaling factors, *L*_1_ and *L*_2_ were defined as:

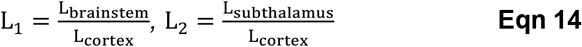

Here L_brainstem_, L_subthalamus_, and L_cortex_ were the anatomical path length from the RVLM to each region and from region RVLM to the cortical measurement site, calculated as Euclidean distance between the three-dimensional (rostrocaudal: rc, mediolateral: ml and dorsoventral: dv) coordinates of the regions using:

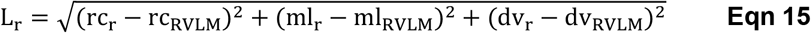

where “r” refers to the region, i.e., brainstem, thalamus, or cortex.

### Statistics

All statistical analyses were performed using custom-written scripts in MATLAB. Data were first tested for normality using the Lilliefors test. For normally distributed datasets, parametric tests such as paired *t*-tests or repeated measures ANOVA were applied as appropriate. For non-normally distributed datasets, non-parametric tests such as Wilcoxon and Friedman’s test were implemented. Statistical significance was defined as *p* < 0.05.

## Data and Software Availability

All the codes are available to access via *GitHub* and data will be made available on request.

## Acknowledgments

We extend our gratitude to Eugene Golanov for discussions and encouragement during the conception of the project, Lauren McElvain for advice on histology, Alex Groissman for assistance with designing the hypoxia chamber, Meenakshi Ashokan, Vineet Augustine, Anna Devor, Patrick Drew, Benjamin Holloway, Clare Howarth and Andy Shih for additional discussions, and Beth Friedman for detailed comments on drafts of the manuscript. This work was supported by the NIH BRAIN initiative grant U19 NS123717 as well as NIH grants U24 EB028942 and R35 NS09726

## Author Contributions

K.C and D.K. conceived the study and designed the experiments, K.C. performed all but the widefield experiments, which were carried out by J.D. P.Y. assisted with the in vivo two-photon measurements and A.F.-Z. assisted with the breathing measurements. K.C. and J. D analyzed data with input from D.K. K. C. and D.K. wrote the manuscript with input from all authors. D.K. attended to the university rules and forms that govern environmental health and safety, including the ethical use of animals as well as the use of chemicals, controlled substances, hazardous substances, lasers, and viruses.

