## Extended data and figures for "A hypoxia-sensitive medullary nucleus modulates cerebral blood flow via a disynaptic pathway to cortex"

### Supplementary Information

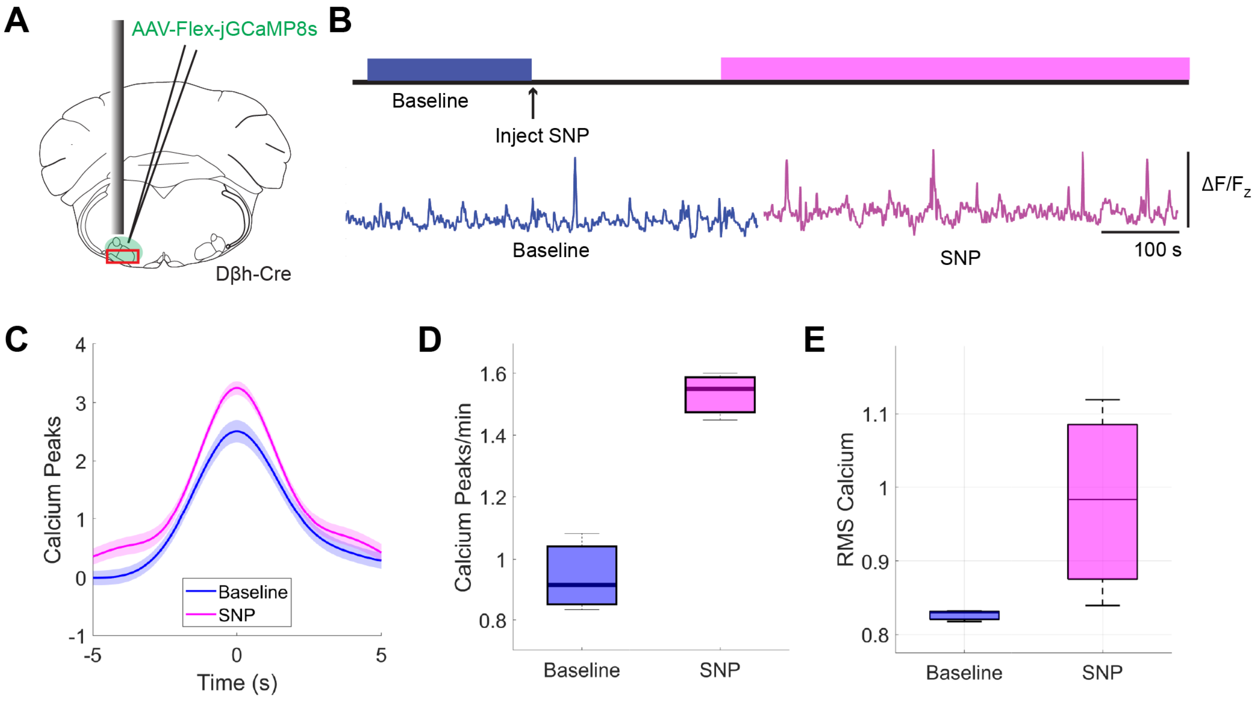


**Extended Figure 1**: Sodium nitroprusside (SNP) increases activity of $RVLM_{D\beta h}$. (A) Injection cartoon showing the location of injection of AAV8-Flex-CAG-jGCaMP8s and fiber implantation in RVLM. (B) Schematic showing timeline of injection and recording with baseline recording period shown in blue and post SNP injection recording in magenta. Corresponding calcium recording for baseline or SNP injection are shown below in blue and magenta, respectively. (C) Peak triggered average showing an increase in peak amplitude under SNP treatment. Error bars are standard error of mean. (D) Average of calcium peaks/min and (E) root mean square (RMS) calcium for the same mice quantified for baseline and SNP recordings.

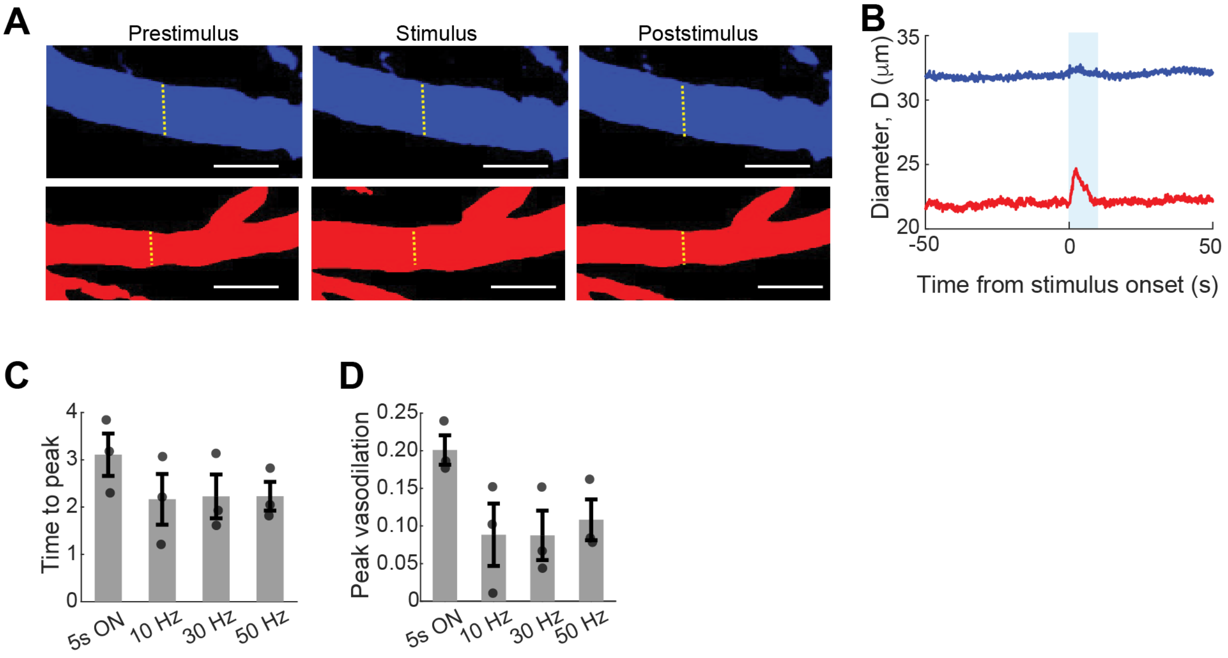


**Extended Figure 2:** Optogenetic activation of dilated arterioles. (A) Example micrograph showing change in cortical venule (blue) and arteriole (red). (B) Trial averaged time series comparing vasodilation amplitude in arteriole and venule (15 and 7 trials, respectively). (C) and (D) shows quantification of time to peak and peak vasodilation in cortical arterioles upon stimulation at continuous 5 s ON pulse, 10, 30 and 50 Hz (3 mice).

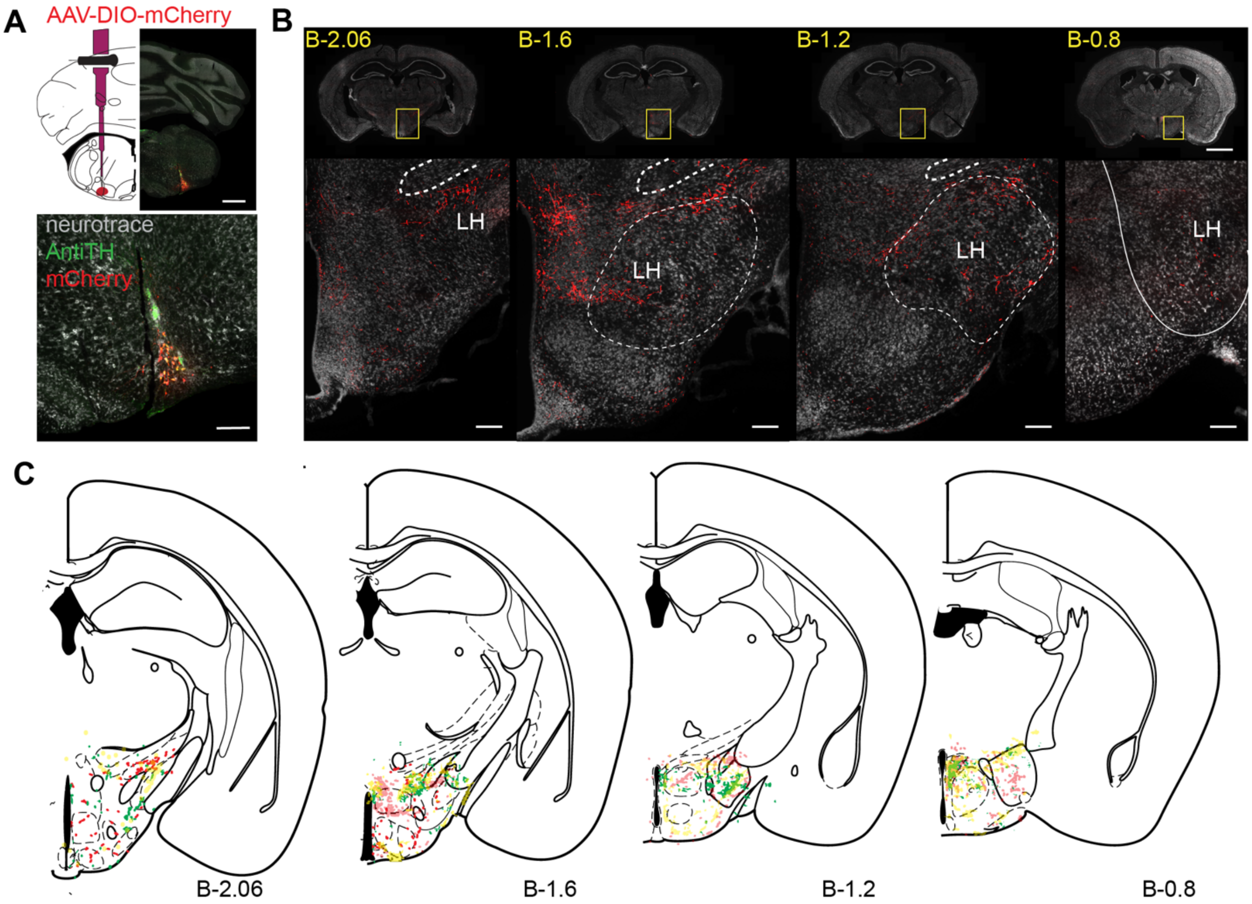


**Extended Figure 3:** Anterograde projections of $RVLM_{D\beta h}$ to lateral hypothalamus (LH). (A) Injection cartoon showing location of injection of AAV8-EF1α-DIO-mCherry and labelled cells. Scale bar is 1 mm in the injection cartoon and 200 µm is subset showing labelled cells. (B) Rostrocaudal coronal sections showing labelled fibers in lateral hypothalamic territories. (C) Rostrocaudal overlays showing lateral hypothalamic labelling across 3 mice.

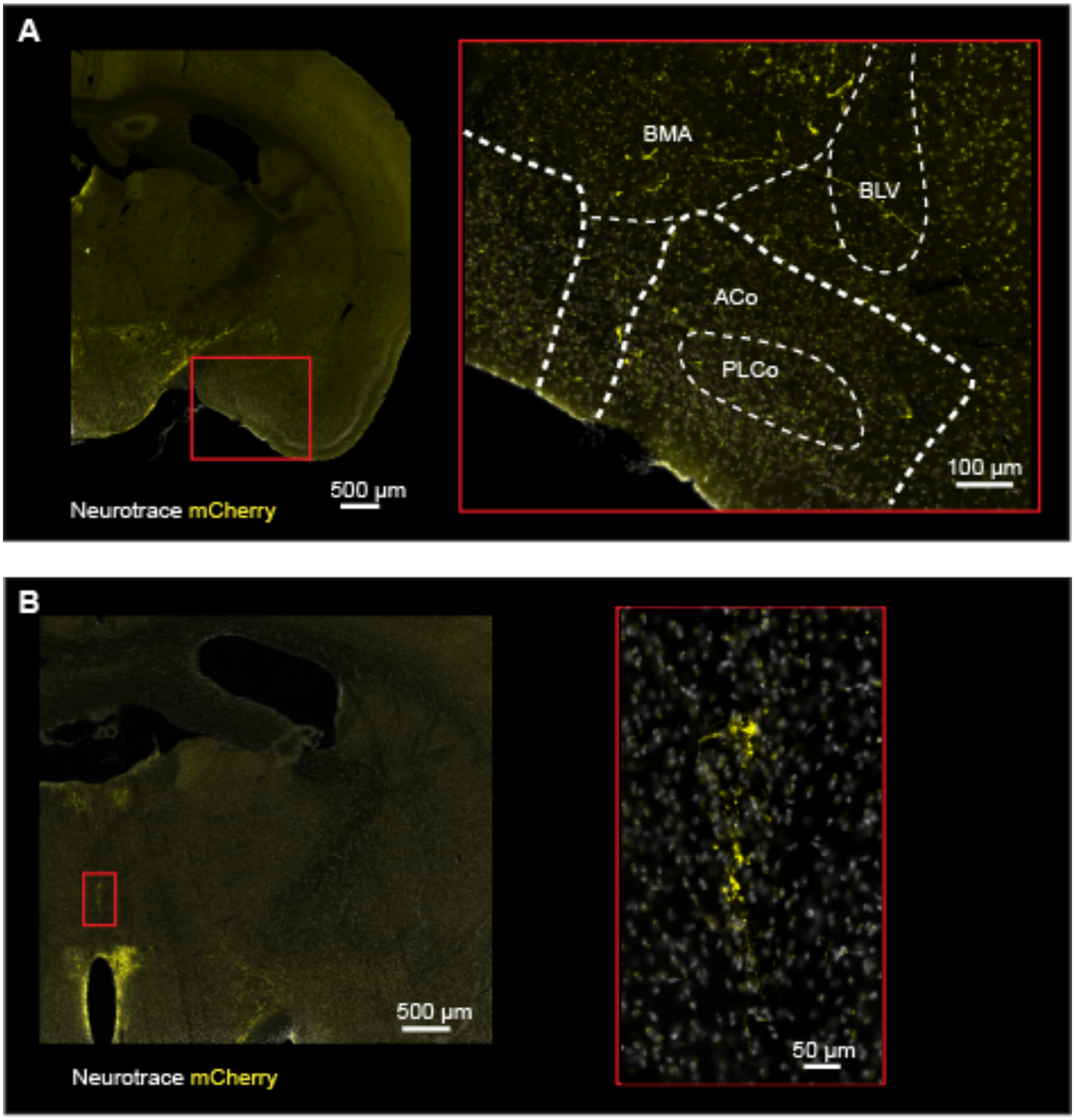


**Extended Figure 4:** Sparse labelling from anterograde tracing from $RVLM_{D\beta h}$ showing labelling in (A) amygdala including basomediolateral amygdala (BMA) and anterior cortical amygdala (ACo) and (B) centromedian (CM) thalamic nucleus. Scale bar: 500 µm (coronal sectional view) and 100 µm.

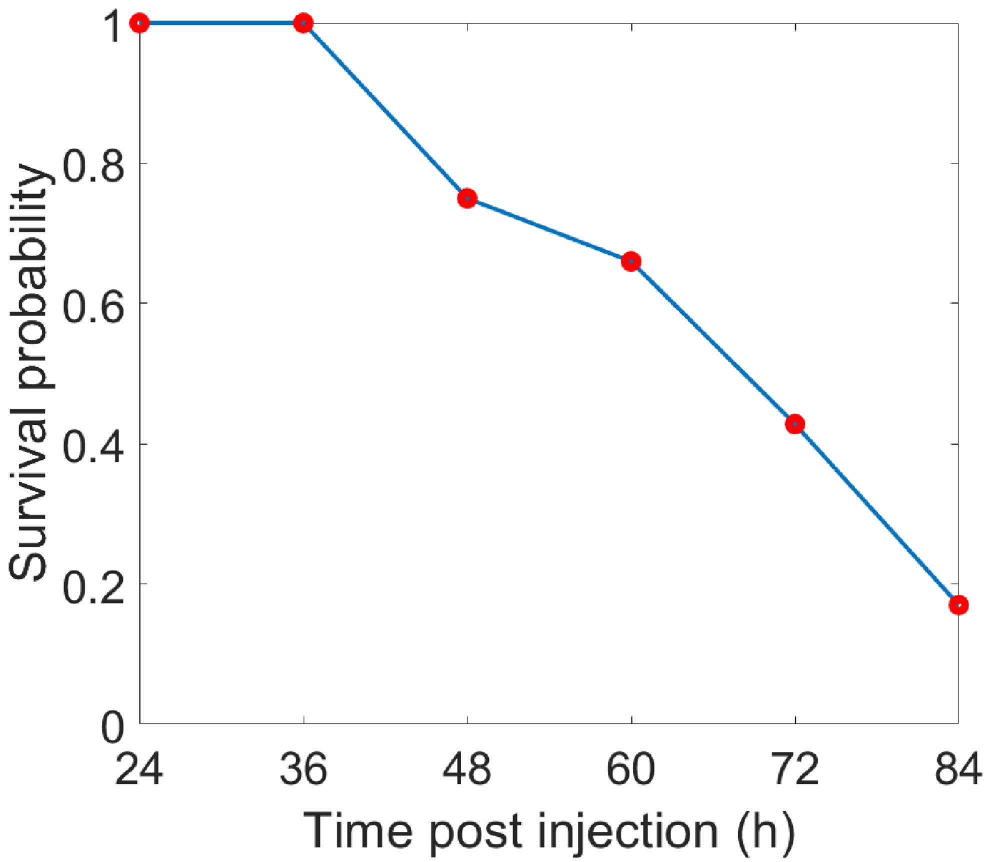


**Extended Figure 5:** Survival plot showing survival probability of mice post injection of HSV-H129-ΔTK-tdtomato.

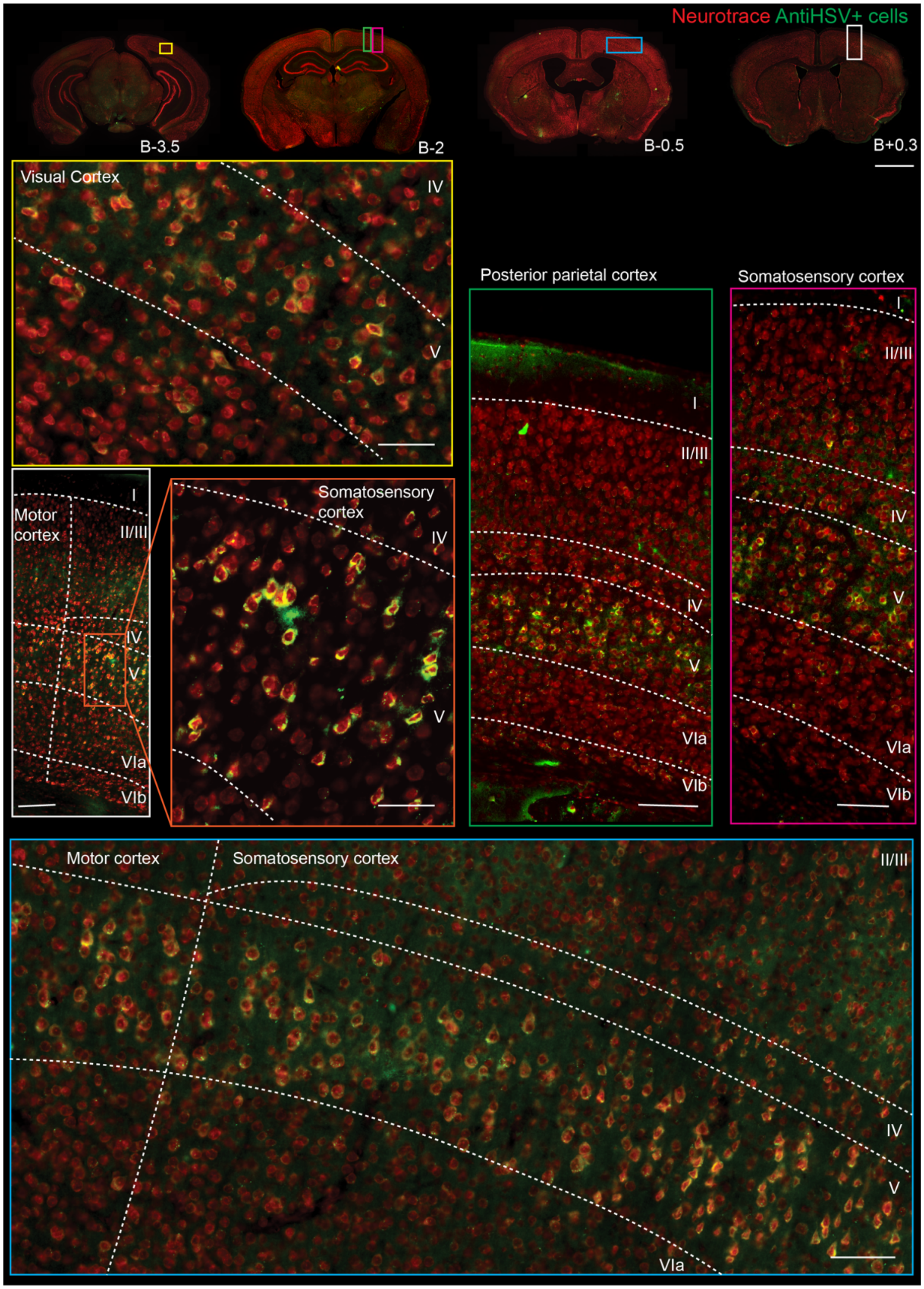


**Extended Figure 6:** Cortical spread of cells labelled from transsynaptic labelling at 72 h post infection. Rostrocaudal coronal sections showing labelling in visual cortex, motor cortex and somatosensory cortex. Most of labelled cells were concentrated around layer V in all cortices with few cells labelled in Layer VIa.

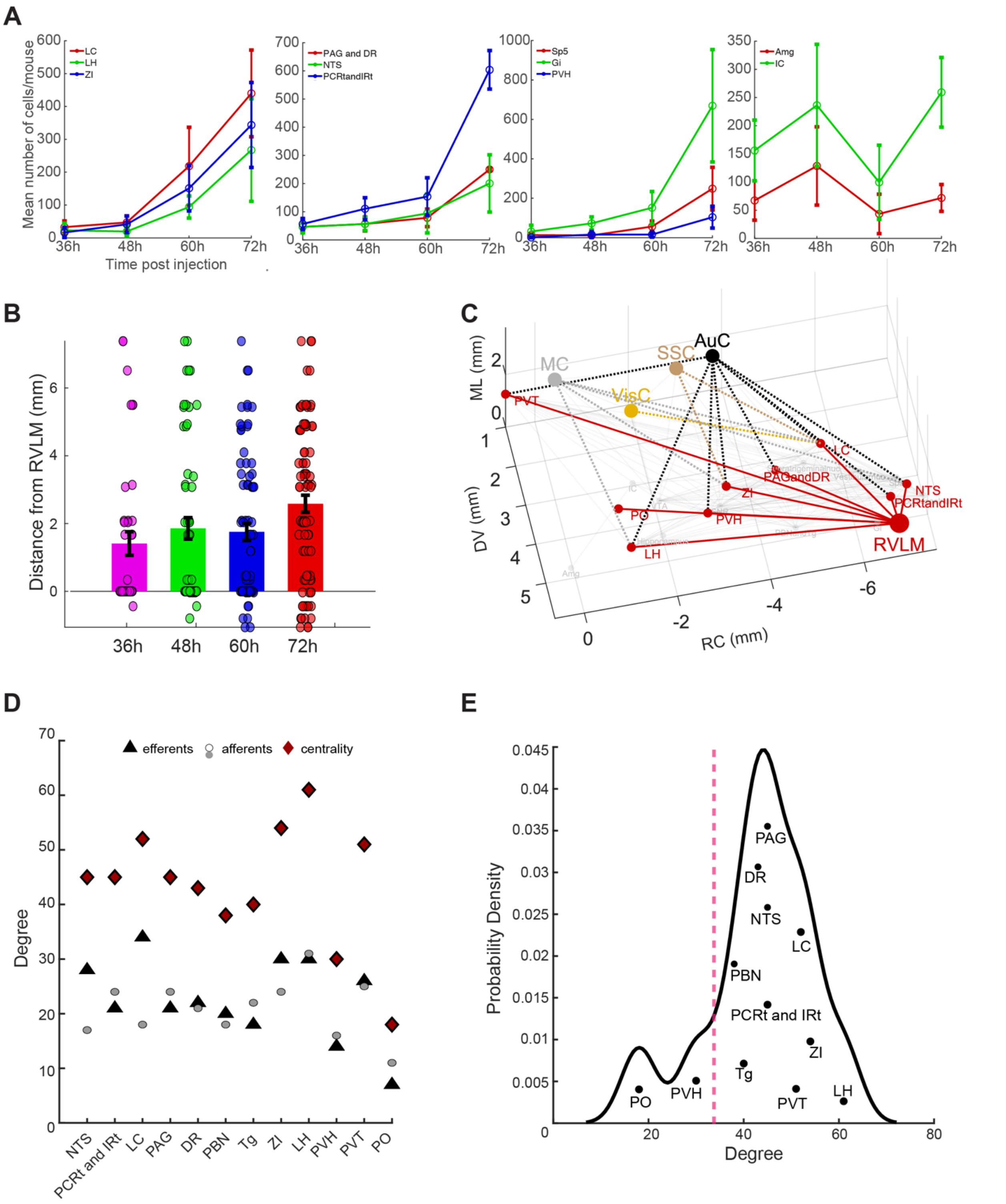


**Extended Figure 7:** Polysynaptic labelling from $RVLM_{D\beta h}$ with HSV-H129-ΔTK-tdTomato shows brain wide labelling of networks with intermediate nodes with high centrality. (A) Mean number of cells across different mice and time points showing examples of set of areas showing similar slope profiles; LH, LC, and ZI; PAG and DR, NTS, and PCRt and IRt; Sp5, Gi, and PVH; and Amg and IC. (B) Mean distance of the cell labelling over time showing an increase in pathlength from RVLM from 36 h to 72 h. (C) Predicted network from the correlation matrix showing primary projections in red (taken from AAV anterograde labelling) and labelling from the primary nodes to the cortical nodes with somatosensory cortex (SSC) in brown, visual cortex (VisC) in yellow, auditory cortex (AuC) in black and motor cortex (MC) in gray. (D) In-, out- degree and centrality (as a measure of total degree of a node = indegrees + outdegrees) of primary nodes from RVLM to primary projections observed in both anterograde non synaptic AAV labelling and transsynaptic HSV labelling. (E) Probability density showing distribution of total degrees of different areas illustrating two ‘humps.’ The red line marks our thresholding for selecting candidates with ‘high’ centrality.

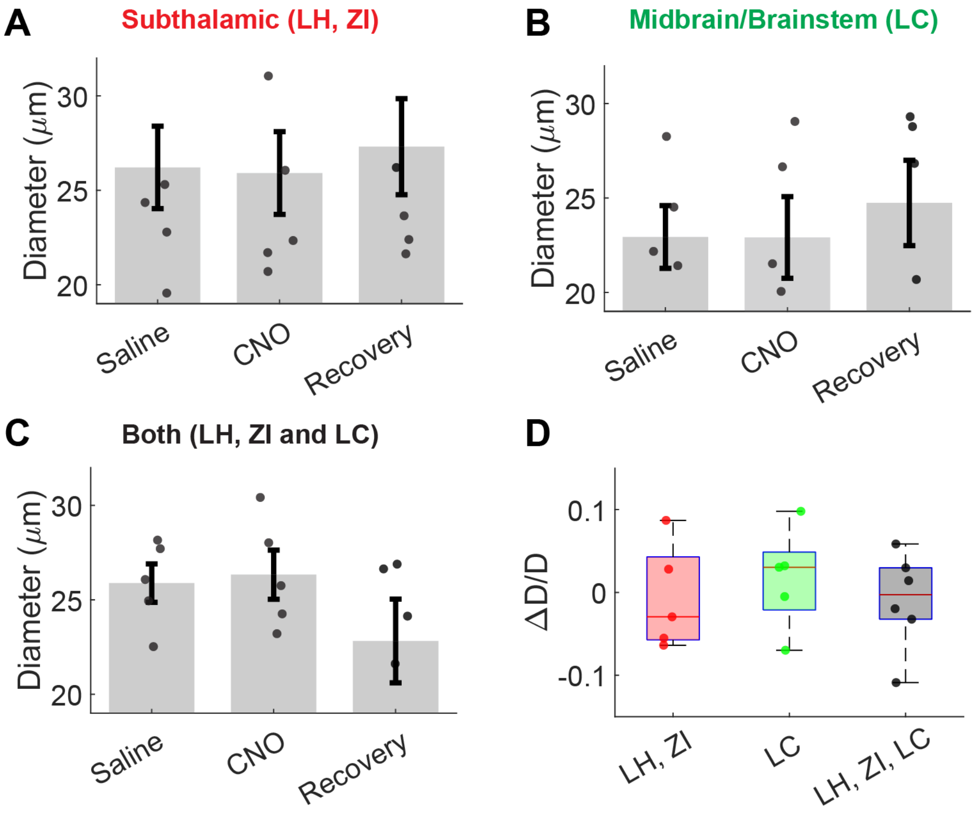


**Extended Figure 8**: Effect of CNO on baseline vessel diameters. Bars showing comparison of absolute values of diameters under saline, CNO and recovery periods with blocking (A) subthalamic nuclei (LH, ZI), (B) midbrain/ brainstem nuclei (LC) and (C) subthalamic and midbrain/brainstem nuclei (LH, ZI and LC). (D) Comparisons of change in diameter with CNO for all three groups showing no significant change in baseline diameters when compared under saline and CNO.

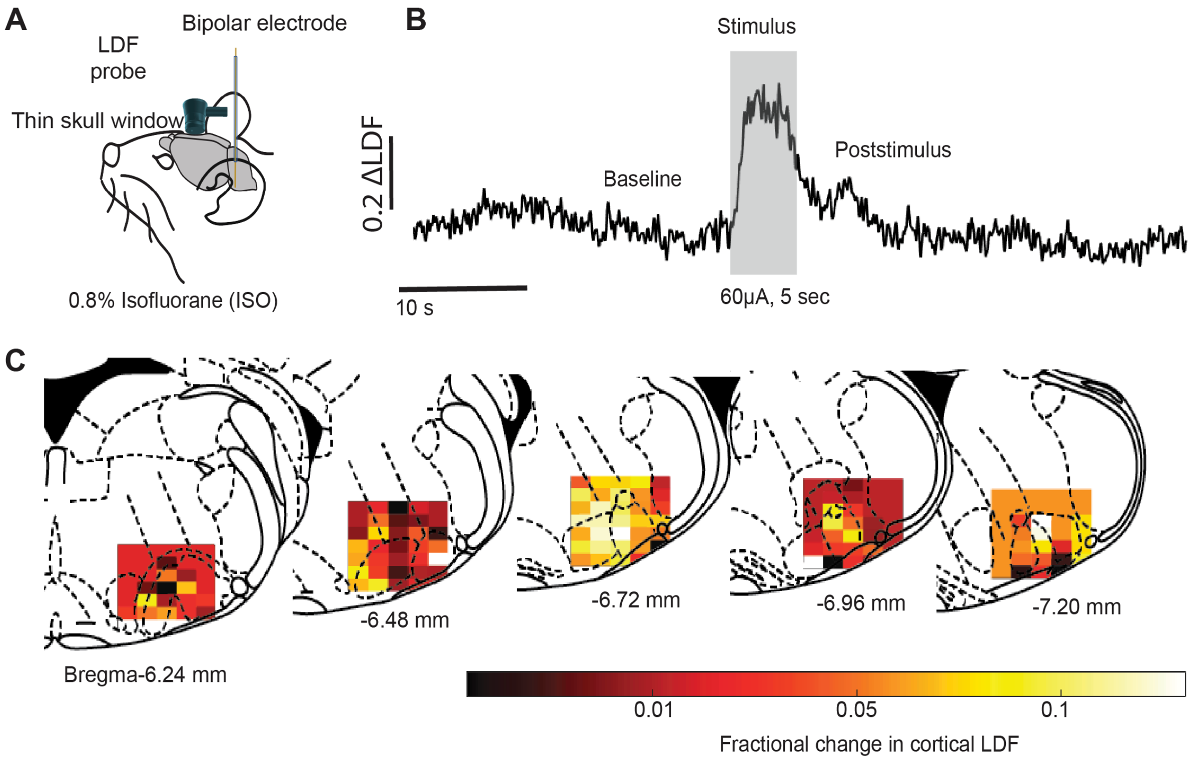


**Extended Figure 9**: Calibration of RVLM coordinates using electrical stimulation in anesthetized mouse. (A) Schematic showing the placement of bipolar electrode for stimulation of RVLM. (B) LDF response to electrical stimulation (60 μA for 5 s) at B - 6.75, 1.25 mediolateral and 4.9 mm dorsoventral from the brain surface. (C) Fractional change in LDF measured from the parietal cortex overlayed onto coordinates of stimulation using anatomical overlays from Paxinos.

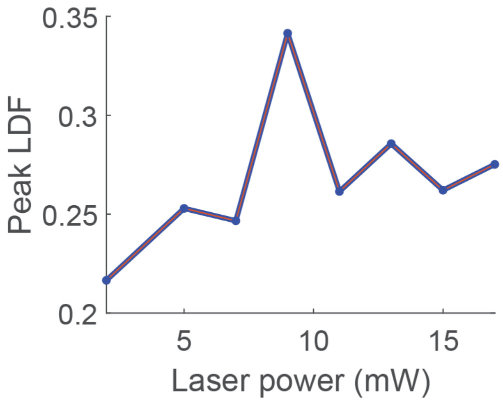


**Extended Figure 10:** Power calibration of optogenetic stimulation to evaluate power values that lead to maximal increase in LDF signal.

**Extended Table 1**: Mean cell slopes of HSV labelling at different time points

| Brain Region | 36 h | 48 h | 60 h | 72 h |
|---|---|---|---|---|
| NTS | 1.292 | 0.833 | 3.146 | 8.875 |
| Hypoglossal nuc | 0.215 | 2.438 | 1.146 | 7.833 |
| PCRt and IRt | 1.576 | 4.479 | 3.625 | 37.479 |
| Sp5 | 0.375 | −0.167 | 3.750 | 16.021 |
| Gi | 0.875 | 3.438 | 6.604 | 43.104 |
| Vestibular nuc | 0.993 | 1.938 | −0.604 | 12.958 |
| LC | 0.910 | 1.188 | 14.271 | 18.458 |
| Supratrigeminal nuc | 0.049 | −0.146 | 2.333 | 9.229 |
| PAG and DR | 1.285 | 0.875 | 1.813 | 14.313 |
| PBN and Tg | 2.597 | 5.438 | 6.188 | 9.250 |
| SNR | 5.042 | 8.104 | 6.563 | 13.980 |
| ZI | 0.458 | 2.063 | 9.146 | 16.042 |
| VTA | 0.194 | 1.479 | 2.479 | 4.042 |
| PVH | 0.042 | 1.188 | 0.021 | 7.333 |
| LH | 0.639 | −0.292 | 6.208 | 14.438 |
| Hippocampus | 0.813 | 1.438 | 45.063 | −0.813 |
| VisC | 1.729 | 14.271 | 63.729 | 24.833 |
| AuC | 0.396 | 11.521 | 7.313 | 48.563 |
| SSC | 2.625 | 45.000 | 51.417 | 58.083 |
| MC | 3.264 | 4.771 | 20.979 | 19.875 |
| PVT | 1.125 | 0.667 | 9.521 | — |
| PO | 0.507 | 1.844 | 2.531 | −2.604 |
| Amg | 1.868 | 5.104 | −7.083 | 2.354 |
| IC | 4.333 | 6.688 | −11.396 | 13.313 |

**Extended Table 2**: Efferents and afferents from primary projections of RVLM

| Region | Efferents | Afferents | Centrality score |
|---|---|---|---|
| NTS[1–6] | RVLM, CVLM, mesencephalic trigeminal nucleus (Me5), locus coeruleus (LC), facial nucleus (FN), reticular nucleus parvicellular (PCRt) and lateral, SpTr, PBN, laterodorsal tegmentum (Tg), paraventricular thalamus (PVT), PVH, paraaqueductal gray (PAG), ventral tegmentum (VTA), substantia niagra reticulata (SNr), lateral hypothalamus (LH), DMH, ventral posteromedial nucleus (VPM), central amygdalar nucleus (CeA), zona incerta (ZI), stria terminalis (ST), ventral pallidum (VP), nucleus | Facial vagal and glossopharangeal nerves, prefrontal cortex (PFC), insular cortex (IC), prelimbic cortex, CeA, ST, PVH, LH, PBN, PAG, Bar, reticular formation parvi and intermediate (PCRt and IRt), DMH, VTA, arcuate nucleus, AP | 46 |

| | | | |
|---|---|---|---|
| | accumbens (NAc), barrington nucleus (Bar), Gigantocellular reticular nucleus (Gi), arcuate nucleus, Subfornical organ (SFO), raphe nucleus, area postrema (AP) | | |
| PCRt and IRt[7–11] | SpTr, mesencephalic trigemenial nucleus, motor trigeminal nucleus, supratrigeminal nucleus, FN, Hypoglossal nucleus, PBN, NTS, inferior olivary nucleus, Gigantocellular reticular nucleus (Gi), nucleus linearis, dorsal reticular nucleus of medulla, ventral reticular nucleus of medulla, NA, DMV, raphe nucleus, RVLM, VRG, spinal cord, cranial nerves, Botzinger complex | Cerebellum, IC, motor cortex (MC), somatosensory cortex (SSC), entorhinal cortex (EC), ST, VP, CeA, PVH, Red nucleus, ZI, LH, SNr, substantia niagra compacta (SNc), substantia inominata, deep mesencephalic nucleus, principal trigeminal nucleus, SpTr, Superior colliculus (SC), PAG, PBN, RVLM, NTS, H2 of Forel | 46 |

| | | | |
|---|---|---|---|
| LC[12–18] | Cerebellum, Hippocampus, LH, PVH, DMN, spinal cord, PBN, laterodorsal Tg, PAG, medial reticular formation, raphe nucleus, NTS, motor trigeminal, hypoglossal nucleus, NA, cochlear nucleus, VTA, SNr, SNc, intralaminar nucleus, posterior thalamic nucleus, VM thalamus, habenula, ST, ZI, lateral genicular nucleus (LGN), globus pallidus (GP), caudate putamen (CP), olfactory cortex (OC), perirhinal cortex, septum (medial and lateral), MC, SSC | NTS, RVLM, nucleus prepositus, Bar, CNA, ST, PFC, CeA, Preoptic nucleus (MPO, PPO, LPO), DMN, PVH, LH, PaF, raphe nucleus, PBN, vestibular nucleus, reticular formation, cerebellum | 52 |
| PAG[19–24] | PVT, DMH, VTA, Septum (medial), ST, RVLM, NTS, NA, intralaminar thalamus, reticular thalamic nucleus, LH, PVH, Superior colliculus, | Spinal cord, NAc, CeA, medial preoptic areas, ST, PFC, MC, VC, LH, SNc, SNr, VTA, ZI, SpTr, PBN, reticular formation (PCRt/IRT), parafasciluar nucleus (PaF), raphe | 45 |

| | | | |
|---|---|---|---|
| | inferior olive, preoptic areas, PBN, spinal trigeminal nucleus, Spinal cord, VRG, fields of forel, parafascicular nucleus. | nucleus, cingulate cortex, septum (lateral), Hippocampus, RVLM, Tg, ventromedial hypothalamus (VMH). | |
| Tegmentum (laterodorsal and pedunculopontine)[25–27] | CC, PFC, septum (medial/diagonal and lateral), preoptic areas, PaF, LH, ZI, laterodorsal and reticular thalamic nuclei, PVT, superior colliculus, inferior olive, pontine reticular nucleus, SNc, VTA, raphe nucleus, hypoglossal nucleus, PBN, NTS | CC, IC, OC, PFC, ST, septum (medial/lateral), CeA, preoptic areas, VP, PaF, ZI, LH, GP, interpeduncular nucleus, lateral mamillary nucleus, SNr, VTA, midbrain central grey, raphe nucleus, LC, PBN, NTS | 39 |
| Dorsal Raphe[28–30] | VP, CP, preoptic area, septum (medial and lateral), venterolateral PAG, VTA, SNc, LH, suprachiasmatic nucleus (SCN), medial CeA, NAc, OC, PFC, EC, IC, CC, hippocampus, rostromedial, | PFC, OC, CC, ST, preoptic area (median and lateral), septum (lateral), VP, LH, tuberomammillary nucleus (TMN), DMH, CeA, VTA, SNc, IP, SNr, PBN, tegmentum (laterodorsal, rosromedial and | 43 |

| | | | |
|---|---|---|---|
| | laterodorsal, pedunculopontine Tg, supraocular nucleus (Su3), reticular formation, RVLM, LC | pedunculopontine), cranial nerve nuclei, RVLM, NTS, NAc | |
| PBN[31–34] | Raphe nucleus (dorsal), edinger Westphal nucleus, PVT, intralaminar thalamus, VM thalamus, PVH, LH, ZI, VP, DMH, arcuate nucleus, preoptic areas (lateral), CeA, ST, RVLM, reticular formation, NTS, Spinal cord (lateral funiculus), IC, PFC | PFC, OC, IC, ST, preoptic area (medial), LH, ZI, DMH, PVH, SNr, raphe nucleus, Edinger Westphal nucleus, Tg (laterodorsal and peduncuplopontine), SpTr, NTS, PAG, VTA, AP | 38 |
| ZI[35–39] | LH, mesencephalic reticular formation, PBN, superior colliculus, red nucleus, Tg (laterodorsal and mesencephalic), PaF, GP, VP, ventromedial thalamus, intralaminar nucleus, subfornical organ, VC, MC, SSC, retrospenial cortex, | Cerebellum, gracile nucleus, SpTr, PAG, mesencephalic reticular nucleus, midbrain raphe, CC, SSC, CeA, PBN, superior colliculus, LGN, LH, pontine reticular nucleus, BLA, rostral ventromedial medulla, VTA, SNr, red nucleus, | 56 |

| | | | |
|---|---|---|---|
| | Spinal cord, PO, basal forebrain, PVT, BMA, LC, cerebellum, VTA, SNr, SNc, PAG, raphe nucleus (dorsal), midbrain reticular nucleus, hippocampus | AC, EC, MC, VC, spinal cord | |
| LH[40–44] | PVH, PO (lateral), ZI, PVT, ST, PAG, VTA, raphe nucleus, Tg (laterodorsal), habenula, PBN, CeA, NTS, A5 catecholamine cell group, RVLM, CC, IC, MC, subfornical organ, VP, superior colliculus, DMV, LC, olfactory nucleus, hippocampus, SNc, reticular formation (medial), arcuate nucleus, ventromedial hypothalamus, anterior hypothalamus | OC, NAc, PFC, IC, olfactory tracts, PO (medial), ST, CP, GP, septum (lateral and medial), CeA, ZI, perifomical region, DMH, PVH, ventral thalamus, fields of Forel, VTA, interpeduncular nucleus, SNr, mesencephalic reticular formation, PAG, LC, PBN, arcuate and mamillary nucleus, supraoptic nucleus, periventricular nucleus of hypothalamus, SCN, NTS, RVLM, Hippocampus | 61 |
| PVH[45,46] | PAG, SNr, raphe nucleus, PBN, LC, reticular nucleus (parvi and lateral), NA, NTS, | Septum (lateral), ventral part of subicular cortex, preoptic area (medial and lateral), subfornical | 30 |

| | | | |
|---|---|---|---|
| | AP, Spinal cord, habenula, LH, PVT, ZI | organ, SCN, LH, ZI, Hippocampus, AP, ST, PVT, CeA, PBN, LC, RVLM, NTS | |
| PVT[47–50] | IC, CC, PFC, EC, perirhinal cortex, dorsal tenia tecta, claustrum, septum (lateral), dorsal striatum, NAc (core and shell), olfactory nucleus, ST, CeA, BMA, BLA, SCN, arcuate nucleus, DMH, ventromedial hypothalamus, PVH, Hippocampus, ventral subiculum, VTA, VP, LH | PBN, PAG, DMN, IC, CC, PFC, Preoptic area (medial), Hippocampus, reticular thalamic nucleus, RVLM, SCN, LH, intergeniculate leaflet, raphe nucleus, NTS, LC, Tg, dorsomedial nucleus, reticular formation, PO, ventromedial hypothalamus, PVH, septum, subfornical organ, CeA | 51 |
| PO[51–53] | LC, raphe nucleus, tuberomammillary nucleus, LH, Tg, PBN, RVLM | Raphe nucleus, RVLM, LC, tuberomammillary nucleus, LH, DMH, infralimbic cortex, PBN, Septum, subiculum, SCN, | 18 |

1. Shapiro, R. E. & Miselis, R. R. The central neural connections of the area postrema of the rat. *J. Comp. Neurol.* **234**, 344–364 (1985).
2. Aicher, S. A., Kurucz, O. S., Reis, D. J. & Milner, T. A. Nucleus tractus solitarius efferent terminals synapse on neurons in the caudal ventrolateral medulla that project to the rostral ventrolateral medulla. *Brain Res.* **693**, 51–63 (1995).
3. Horst, G. J. T. & Streefland, C. Ascending Projections of the Solitary Tract Nucleus. in *Nucleus of the Solitary Tract* (CRC Press, 1994).
4. Gasparini, S., Howland, J. M., Thatcher, A. J. & Geerling, J. C. Central afferents to the nucleus of the solitary tract in rats and mice. *J. Comp. Neurol.* **528**, 2708–2728 (2020).
5. Sequeira, S. M., Geerling, J. C. & Loewy, A. D. Local inputs to aldosterone-sensitive neurons of the nucleus tractus solitarius. *Neuroscience* **141**, 1995–2005 (2006).
6. Holt, M. K. The ins and outs of the caudal nucleus of the solitary tract: An overview of cellular populations and anatomical connections. *J. Neuroendocrinol.* **34**, e13132 (2022).
7. Almeida, A., Cobos, A., Tavares, I. & Lima, D. Brain afferents to the medullary dorsal reticular nucleus: a retrograde and anterograde tracing study in the rat. *Eur. J. Neurosci.* **16**, 81–95 (2002).
8. Ter Horst, G. J., Copray, J. C. V. M., Liem, R. S. B. & Van Willigen, J. D. Projections from the rostral parvocellular reticular formation to pontine and medullary nuclei in the rat: Involvement in autonomic regulation and orofacial motor control. *Neuroscience* **40**, 735–758 (1991).

51. Sherin, J. E., Elmquist, J. K., Torrealba, F. & Saper, C. B. Innervation of histaminergic tuberomammillary neurons by GABAergic and galaninergic neurons in

the ventrolateral preoptic nucleus of the rat. *J. Neurosci. Off. J. Soc. Neurosci.* **18**, 4705–4721 (1998).

52. Chou, T. C. *et al.* Afferents to the Ventrolateral Preoptic Nucleus. *J. Neurosci.* **22**, 977–990 (2002).

53. Arrigoni, E. & Fuller, P. M. The Sleep-Promoting Ventrolateral Preoptic Nucleus: What Have We Learned over the Past 25 Years? *Int. J. Mol. Sci.* **23**, 2905 (2022).
